# Extensive profiling of histidine-containing dipeptides reveals species- and tissue-specific distribution and metabolism in mice, rats and humans

**DOI:** 10.1101/2023.02.16.528841

**Authors:** Thibaux Van der Stede, Jan Spaas, Sarah de Jager, Jana De Brandt, Camilla Hansen, Jan Stautemas, Bjarne Vercammen, Siegrid De Baere, Siska Croubels, Charles-Henri Van Assche, Berta Cillero Pastor, Michiel Vandenbosch, Ruud Van Thienen, Kenneth Verboven, Dominique Hansen, Thierry Bové, Bruno Lapauw, Charles Van Praet, Karel Decaestecker, Bart Vanaudenaerde, Bert O Eijnde, Lasse Gliemann, Ylva Hellsten, Wim Derave

**Affiliations:** Department of Movement and Sports Sciences, Ghent University, Belgium; Department of Nutrition, Exercise and Sports, Copenhagen University, Denmark; University MS Center (UMSC) Hasselt – Pelt, Belgium; BIOMED Biomedical Research Institute, Hasselt University, Belgium; REVAL Rehabilitation Research Center, Hasselt University, Belgium; Department of Pathobiology, Pharmacology and Zoological Medicine, Ghent University, Belgium; The Maastricht MultiModal Molecular Imaging (M4I) institute, Maastricht University, The Netherlands; Heart Center Hasselt, Jessa Hospital Hasselt, Belgium; Department of Cardiac Surgery, Ghent University Hospital, Belgium; Department of Endocrinology, Ghent University Hospital, Belgium; Department of Urology, Ghent University Hospital, Belgium; Department of Human Structure and Repair, Ghent University, Belgium; Department of Chronic Diseases and Metabolism, KU Leuven, Belgium; SMRC Sports Medical Research Center, BIOMED Biomedical Research Institute, Hasselt University, Belgium; Division of Sport Science, Stellenbosch University, South Africa

**Keywords:** histidine-containing dipeptides, central nervous system, muscle, exercise

## Abstract

Histidine-containing dipeptides (HCDs) are pleiotropic homeostatic molecules linked to inflammatory, metabolic and neurological diseases, as well as exercise performance. Using a sensitive UHPLC-MS/MS approach and an optimized quantification method, we performed a systematic and extensive profiling of HCDs in the mouse, rat, and human body (in n=26, n=25, n=19 tissues, respectively). Our data show that tissue HCD levels are uniquely regulated by carnosine synthase (CARNS1), an enzyme that was preferentially expressed by fast-twitch skeletal muscle fibers and brain oligodendrocytes. Cardiac HCD levels are remarkably low compared to other excitable tissues. Carnosine is unstable in human plasma, but is preferentially transported within red blood cells in humans but not rodents. The low abundant carnosine analog N-acetylcarnosine is the most stable plasma HCD, and is enriched in human skeletal muscles. Here, N-acetylcarnosine is continuously secreted into the circulation, which is further induced by acute exercise in a myokine-like fashion. Collectively, we provide a novel basis to unravel tissue-specific, paracrine, and endocrine roles of HCDs in human health and disease.

**Significance statement:** By extensively profiling the pluripotent histidine-containing dipeptides across three species, we generated many new insights into species- and tissue-specific histidine-containing dipeptide metabolism. For instance, the only stable analog that is specific for the human circulation (N-acetylcarnosine) is continuously released from muscle tissue and is positively regulated by physical exercise. The great number of analyses and experiments involving humans establishes great translational value of the findings. These new data open exciting opportunities to study histidine-containing dipeptide metabolism, including paracrine and/or endocrine signaling of these dipeptides, possibly contributing to the potent health-promoting exercise effects.

## Introduction

Carnosine synthase (CARNS1) is presumably the only enzyme in animals capable of synthesizing an abundant class of endogenous dipeptides. The enzyme links L-histidine to either β-alanine or γ-aminobutyric acid (GABA), respectively rendering carnosine or homocarnosine. These parent dipeptides and their methylated (anserine and balenine) and acetylated (N-acetylcarnosine) analogs are collectively called the histidine-containing dipeptides (HCDs).

Since the initial discovery in 1900 by Vladimir Gulevich (*1*), carnosine and the other HCDs have been linked to various physiological functions, mostly serving to preserve redox status and cellular homeostasis (for a full overview, see (*2*)). The most relevant biochemical properties for their functions relate to proton buffering, metal chelation and antioxidant capacity, which further translates to protection against advanced glycation and lipoxidation end products (*3, 4*). The physiological importance of tissue HCD content is underscored by an extensive body of research ranging from enhancement of exercise performance (*5*) to treatment of cardiometabolic (*6, 7*) or neurological diseases (*8*) in rodents. Major differences between animal and human HCD metabolism may be present however, given that high carnosinase (CN1) activity in human, but not rodent, plasma results in rapid degradation of carnosine (*9, 10*).

Nevertheless, even more than 120 years after the initial discovery of carnosine and 10 years after the molecular identification of CARNS1 (*11*), there remains a lack of basic understanding of HCD synthesis, distribution, and metabolism throughout the animal and especially the human body. It is thought that HCDs are primarily expressed in excitable tissues such as skeletal and cardiac muscle and the central nervous system (CNS), but current literature mostly consists of scattered observations focussing on a limited number of tissues or species. Information on cardiac levels is sparse, although HCDs could play an important role in cardiomyocyte homeostasis (*12*). Furthermore, there is unclarity regarding the synthesis and physiological role of HCDs in kidney, lung, liver, and other non-excitable tissues. A first profiling of HCDs in rat tissues from Aldini *et al*. (*13*) did not detect HCDs in non-excitable tissues, although this and other previous endeavours were potentially limited from lower detection sensitivity compared to the currently available technology. For example, the low abundant HCDs anserine, balenine and N-acetylcarnosine have never been extensively characterized in animal or human tissues.

Here, we have performed the first systematic profiling of the five main HCDs, combined with determination of CARNS1 expression levels, in the mouse, rat and human body. Various human tissue samples were collected from live donors, except for post-mortem collected brain regions. We uncovered profound differences in HCD distribution and metabolism between tissues and species. For instance, we demonstrate that humans have a unique way of circulating HCDs and releasing it from carnosine-synthesizing tissues such as skeletal muscle.

## Results

### CARNS1 is the unique and rate-limiting enzyme for HCD synthesis

Using whole-body *Carns1*-knockout (KO) mice, we aimed to investigate whether CARNS1 deficiency results in a complete lack of endogenous (homo)carnosine and their derivatives in a variety of tissues, which would imply that CARNS1 is the unique and rate-limiting enzyme for HCD synthesis. CARNS1 was successfully knocked out at the gene (**Fig 1A**) and protein (100 kDa, **Fig 1B**) level, thereby also validating our Western blot antibody for specifically detecting the CARNS1 protein. As described previously, *Carns1*-KO mice displayed normal growth and survival (*14*). The deletion of *Carns1* led to an absence of carnosine and homocarnosine in all investigated tissues (**Fig 1C)**. Similarly, these mice were devoid of the carnosine-derived analogs anserine, balenine and N-acetylcarnosine (of which only anserine is consistently present in mouse tissues, cfr. infra, **Fig 1C**).

**Figure 1.**
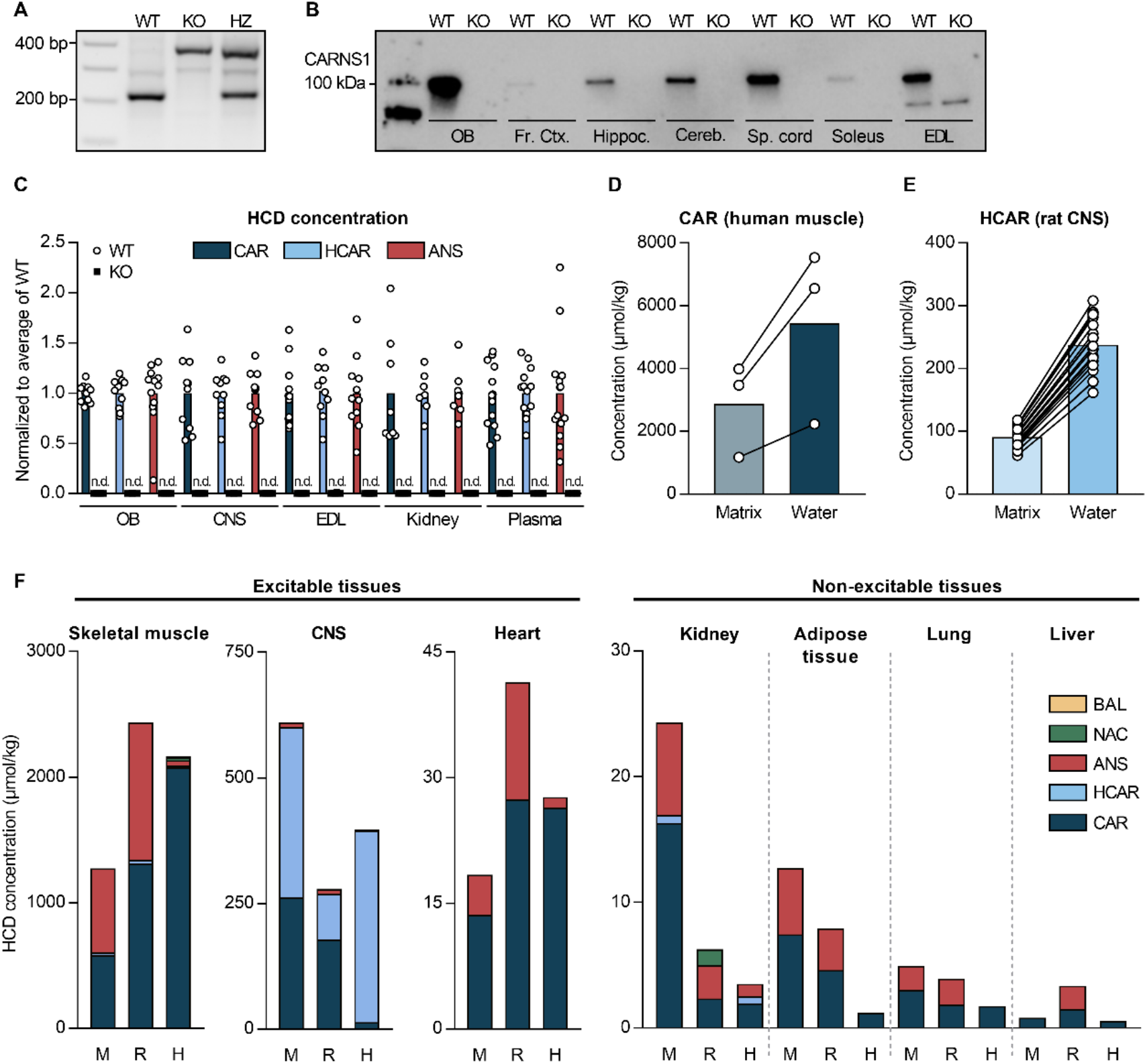
Species- and tissue-specific distribution of histidine-containing dipeptides. **(A)** PCR gel and **(B)** Western blot showing the successful knockout of the *Carns1* gene and the absence of CARNS1 protein in mice. **(C)** HCD measurements by UHPLC-MS/MS showing HCDs are absent from various tissues of *Carns1*-KO compared to WT mice. **(D)** Carnosine and **(E)** homocarnosine measured by UHPLC-MS/MS, followed by quantification based on a standard calibration curve prepared in water *vs. Carns1*-KO tissue matrix. **(F)** HCD measurements by UHPLC-MS/MS in skeletal muscle, CNS, heart, kidney, adipose tissue, lung, and liver tissue from mice (M), rats (R) and humans (H). Values were averaged if more than 1 type of the respective tissue was present (e.g. soleus and EDL for rodent muscle). ANS, anserine; BAL, balenine; CAR, carnosine; Cereb., cerebellum; CNS, central nervous system; EDL, extensor digitorum longus; Fr. ctx., frontal cortex; HCAR, homocarnosine; Hippoc., hippocampus; HZ, heterozygous; KO, knockout; NAC, N-acetylcarnosine; OB, olfactory bulb; Sp. cord spinal cord; WT, wild type.

In addition, tissue from *Carns1*-KO mice was used to optimise our quantitative UHPLC-MS/MS-based detection of HCDs. Concentration levels in muscle and brain were compared using matrix-matched standard calibration curve preparation in *Carns1*-KO tissue matrix (i.e. muscle or brain homogenates) and water (as current standard practice in HCD research). In water, HCD levels were overestimated by ~2 to 3-fold (**Fig 1D-E**), indicating the importance of utilizing a corresponding blank tissue matrix for HCD quantification. This approach was used for all further analyses in this paper (except human cerebrospinal fluid), rendering the HCD quantifications more accurate than previously reported.

### HCDs are not excitable tissue-specific metabolites

Using our sensitive UHPLC-MS/MS method, we performed a systematic profiling of HCDs in various tissues of the mouse, rat, and human body (**Fig 1F**, **Table S1**). Carnosine was the only HCD present in all studied tissues across the three species. Skeletal muscles contained the largest amounts of HCDs, followed by the CNS, up to the millimolar range. However, not all excitable tissue contained large amounts of HCDs, since unexpectedly low HCD levels were observed in the heart of all three species (50-100 times lower than skeletal muscles). These low HCD levels in cardiac tissue better reflect those measured in non-excitable tissues. Besides skeletal muscle, CNS, heart, kidney, adipose, lung and liver tissue (presented in **Fig 1F**), we also found low levels of HCDs in the mouse and rat stomach wall, gallbladder, pancreas, small intestine, colon, thymus, spleen, and eye (**Table S1**).

### CARNS1 content drives fiber type-related differences in HCD content in skeletal muscle

We next aimed to profile the HCD content in skeletal muscle in more detail, with a focus on potential fiber type-specific differences. In mice and rats, we determined the HCD content in soleus (more oxidative, slow-twitch) and extensor digitorum longus (EDL; more glycolytic, fast-twitch) muscles. Our results confirmed previous reports (*15–17*) that carnosine and anserine content is higher in EDL muscle (**Fig 2A**). Also the homocarnosine content was ~2.5-fold higher in EDL compared to soleus in both mice and rats (**Fig 2B**). To get more insights if CARNS1 content (i.e. HCD production) is the main driver for the fiber type-related differences, we first explored a publicly available human muscle fiber type-specific RNAseq dataset (*18*). These data show a ~2-fold higher *CARNS1* mRNA content in type IIa *vs*. type I fibers (**Fig 2C**). We next determined CARNS1 protein levels in the soleus and EDL muscles from the mice and rats (**Fig 2D-E**). CARNS1 levels were indeed higher in EDL muscles, with a larger difference between soleus and EDL muscles in mice (6.6-fold) than in rats (1.4-fold). To translate mRNA differences in human muscle to the protein level, we used Western blotting on pools of pre-classified type I or type IIa fibers (**Fig S1**). This revealed a ~2-fold higher CARNS1 content in type IIa fibers (**Fig 2D-E**), consistent with findings at the mRNA level. These results suggest that CARNS1 expression is the main driver regulating the clear fiber type-related differences in HCD content across the three species.

**Figure 2.**
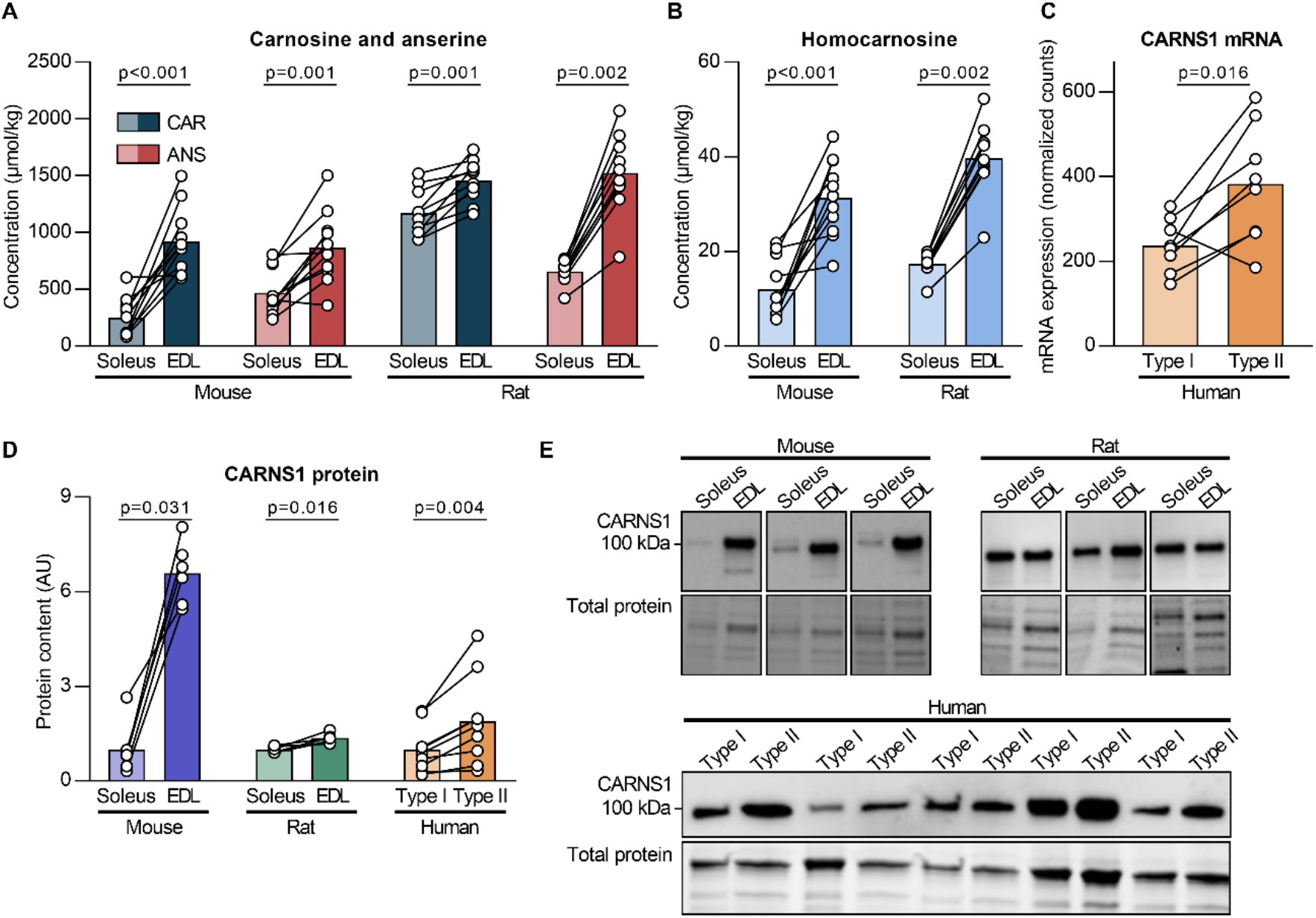
Muscle and muscle fiber type-specific differences in histidine-containing dipeptides and CARNS1. **(A)** Carnosine and anserine and (**B**) measurements by UHPLC-MS/MS in soleus and EDL muscles from mice and rats. **(C)** Human muscle fiber type-specific calculation of *CARNS1* mRNA based on a previously published dataset. **(D)** Protein levels of CARNS1 determined by Western blot in soleus and EDL muscle from mice and rats, and human type I and type II fiber pools. **(E)** Representative Western blot and loading controls. ANS anserine; CAR, carnosine; EDL, extensor digitorum longus.

### Homocarnosine is the dominant HCD in the CNS, except for the olfactory bulb of rodents

Apart from skeletal muscle, HCDs are also highly present in the CNS. By analyzing seven different regions from the mouse, rat and human CNS, we could confirm that the highest levels of carnosine are found in the olfactory bulb in rodents, reaching concentrations of ~1200 μmol/kg tissue, which is similar to or even higher than skeletal muscle carnosine levels (**Fig 3A, Table S1)**. In contrast, the human olfactory bulb contained approximately 15 times less carnosine (~80 μmol/kg). In mice and rats, the olfactory bulb was the only CNS region containing more carnosine than homocarnosine, whilst in humans all regions contained more homocarnosine than carnosine. Similar to our findings in human CNS tissue, homocarnosine was also abundantly present in human cerebrospinal fluid (**Fig 3B**).

**Figure 3.**
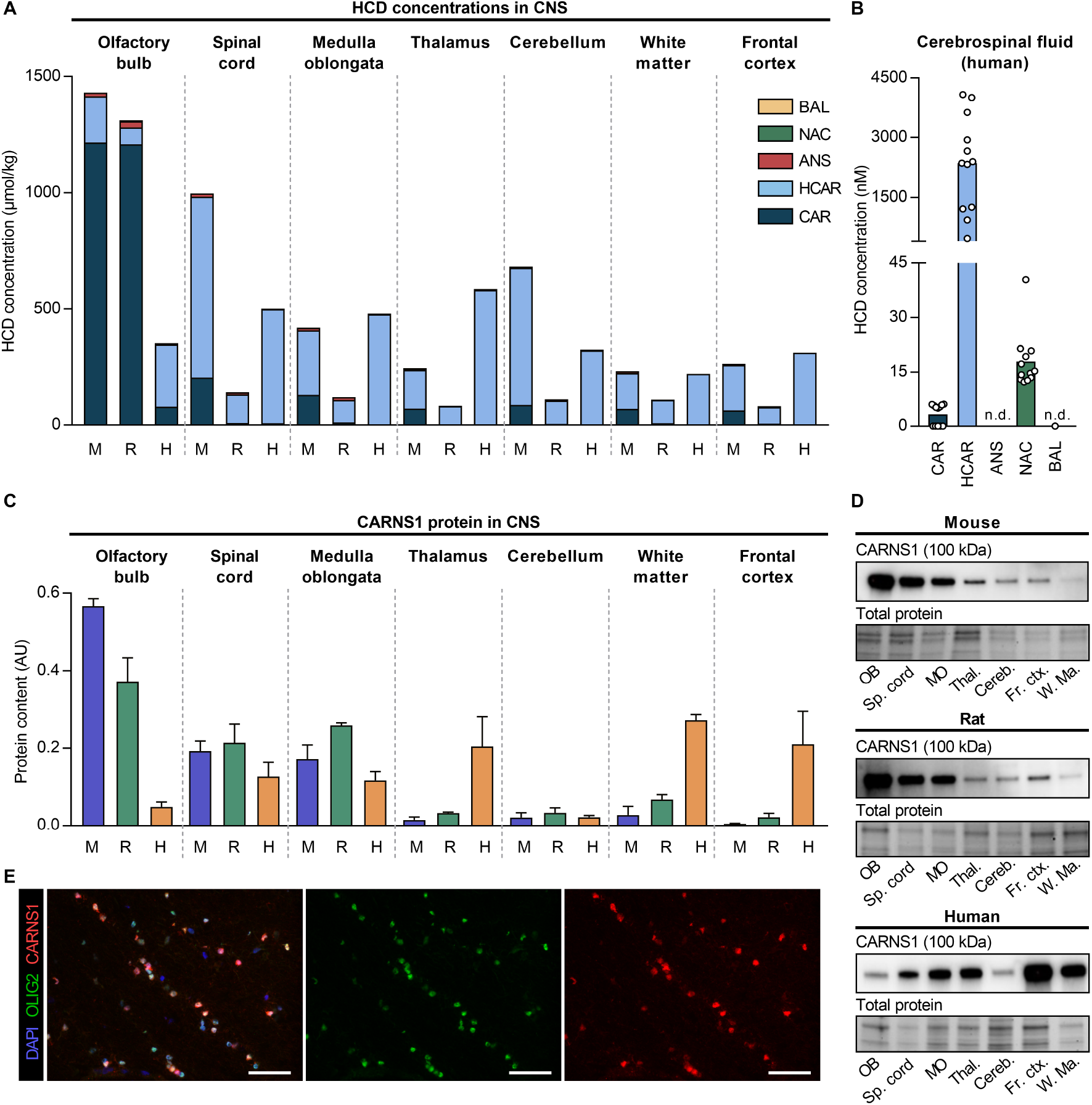
Region-specific levels of histidine-containing dipeptides and CARNS1 in the mouse, rat and human central nervous system. **(A)** HCD measurements by UHPLC-MS/MS in seven different central nervous system regions from mice (M), rats (R) and humans (H). **(B)** HCD measurements by UHPLC-MS/MS in human cerebrospinal fluid. **(C)** Protein levels of CARNS1 determined by Western blot in seven different central nervous system regions from mice, rats and humans. Data are mean ± SD. **(D)** Representative Western blot and loading controls. **(E)** Immunohistochemical detection of CARNS1 and OLIG2 in human white matter. In panels (A), (C) and (D), spinal cord tissue is from the cervical region, mouse white matter is from the corpus callosum, and human frontal cortex is from the superior frontal gyrus. ANS anserine; AU, arbitrary units; BAL, balenine; CAR, carnosine; Cereb., cerebellum; CNS, central nervous system; Fr. ctx., frontal cortex; HCAR, homocarnosine; MO, medulla oblongata; NAC, N-acetylcarnosine; OB, olfactory bulb; Sp. cord spinal cord; Thal., thalamus; W. Ma., white matter. Scale bars are 50 μm (E).

CARNS1 protein levels also showed considerable variability between CNS regions (**Fig 3C-D**). Mice and rats exhibited high expression in olfactory bulb, as well as the spinal cord and medulla oblongata, but not the rest of the CNS. In human tissues, we found lower CARNS1 levels in the olfactory bulb, but instead observed greater amounts in the white matter, thalamus, and frontal cortex.

Immunofluorescence was used to further study the localisation of CARNS1 in the CNS. We chose the region exhibiting the greatest CARNS1 protein levels among rodents (mouse olfactory bulb) and humans (white matter). Double-labeling of CARNS1 and OLIG2, an oligodendrocyte lineage marker, revealed that CARNS1 resides in oligodendrocytes of human white matter (**Fig 3E**). Cell markers for microglia (CD68, **Fig S2A**), astrocytes (GFAP, **Fig S2B**) and neurons/axons (NF-H, **Fig S2C**) did not co-localise with CARNS1. In the mouse olfactory bulb, CARNS1 appeared in spherical structures near the surface of the olfactory bulb, i.e. the glomeruli, where olfactory nerve terminals form synapses with dendrites from projection neurons that carry signals into the brain (**Fig 4A**). A similar spatial distribution was present for carnosine, as mass spectrometry imaging detected higher levels at the border compared to the center of the mouse olfactory bulb (**Fig 4B**). In contrast, homocarnosine displayed a more dispersed distribution throughout the mouse CNS, as well as an overall anterior-to-posterior gradient with greater amounts of homocarnosine in the midbrain, hindbrain, cerebellum, and spinal cord (**Fig 4C**). As expected, (homo)carnosine could not be detected in *Carns1*-KO mice (**Fig 4B-D**).

**Figure 4.**
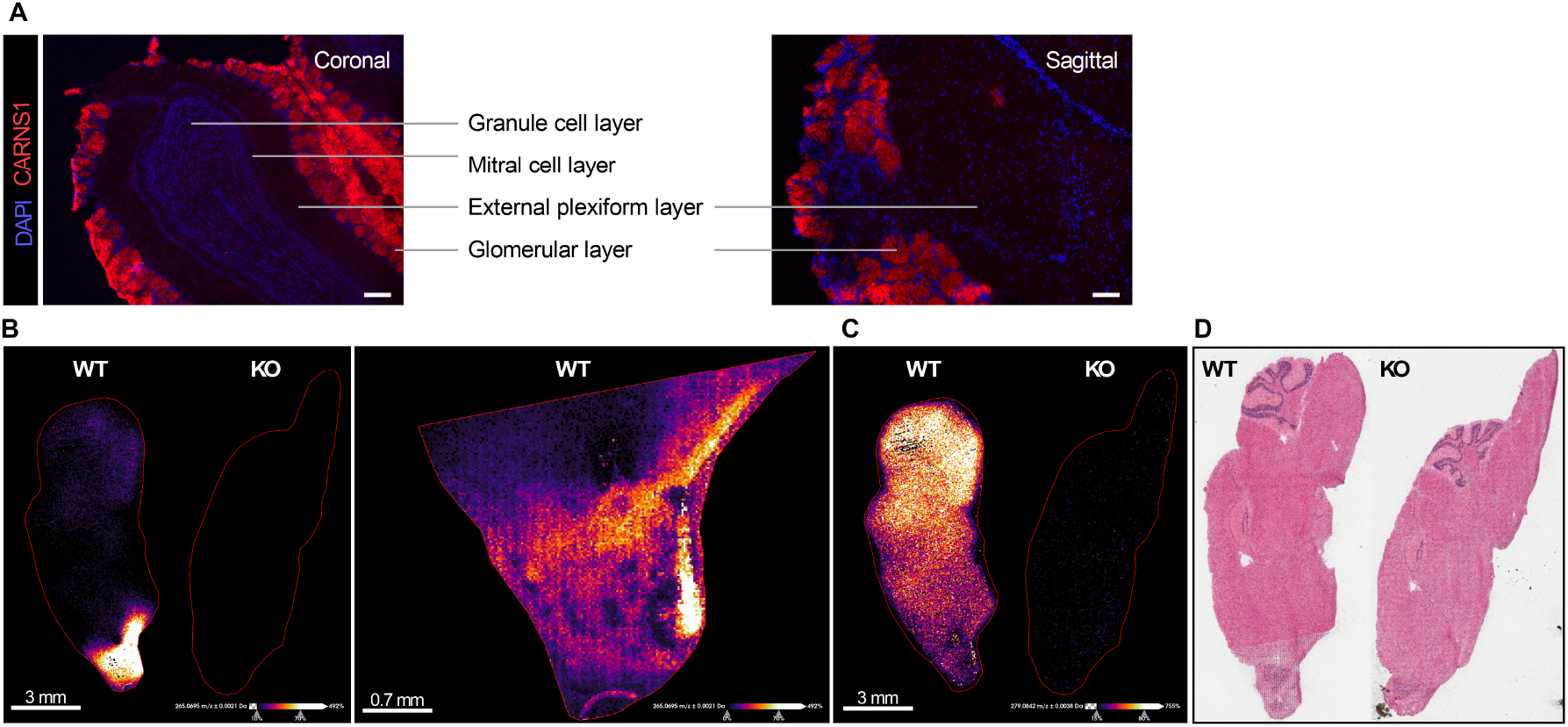
Spatial localisation of CARNS1 and (homo)carnosine within the mouse central nervous system. **(A)** Immunohistochemical detection of CARNS1 in mouse olfactory bulb. Scale bars are 100 μm. **(B)** Mass spectrometry imaging (MALDI-MSI) of carnosine in whole mouse brain (50 μm spatial resolution) or olfactory bulb only (20 μm spatial resolution). WT and *Carns1*-KO mouse. **(C)** Mass spectrometry imaging (MALDI-MSI) of homocarnosine in whole mouse brain (50 μm spatial resolution). WT and *Carns1*-KO mouse. **(D)** H&E staining of the representative WT and *Carns1*-KO mouse brains. KO knockout; WT, wild type.

### CARNS1 expression scales with tissue HCD levels on a whole-body level

To investigate if tissue CARNS1 expression closely relates to tissue HCD levels, CARNS1 protein levels (Western blot) were plotted against HCD concentrations (UHPLC-MS/MS). If the tissue CARNS1 level is the main determinant of tissue HCD levels, a linear relation between both variables is expected. In all non-excitable tissues, no CARNS1 could be detected by Western blot (**Fig 5A-C**). HCD levels in these tissues probably reflect transmembrane HCD uptake. On a whole-body level, CARNS1 scaled with HCD levels in mice (**Fig 5A**), rats (**Fig 5B)** and humans (**Fig 5C**). However, when comparing within organs, more CARNS1 was not always directly linked to a higher HCD concentration. In the human CNS, for example, CARNS1 was 13-fold higher in white matter than in the cerebellum, but both tissues had similar HCD levels. This indicates that although CARNS1 is the only enzyme responsible for HCD synthesis, other factors, such as exchange of HCDs between organs or a high HCD turnover rate (synthesis/degradation), could also influence intracellular HCD levels.

**Figure 5.**
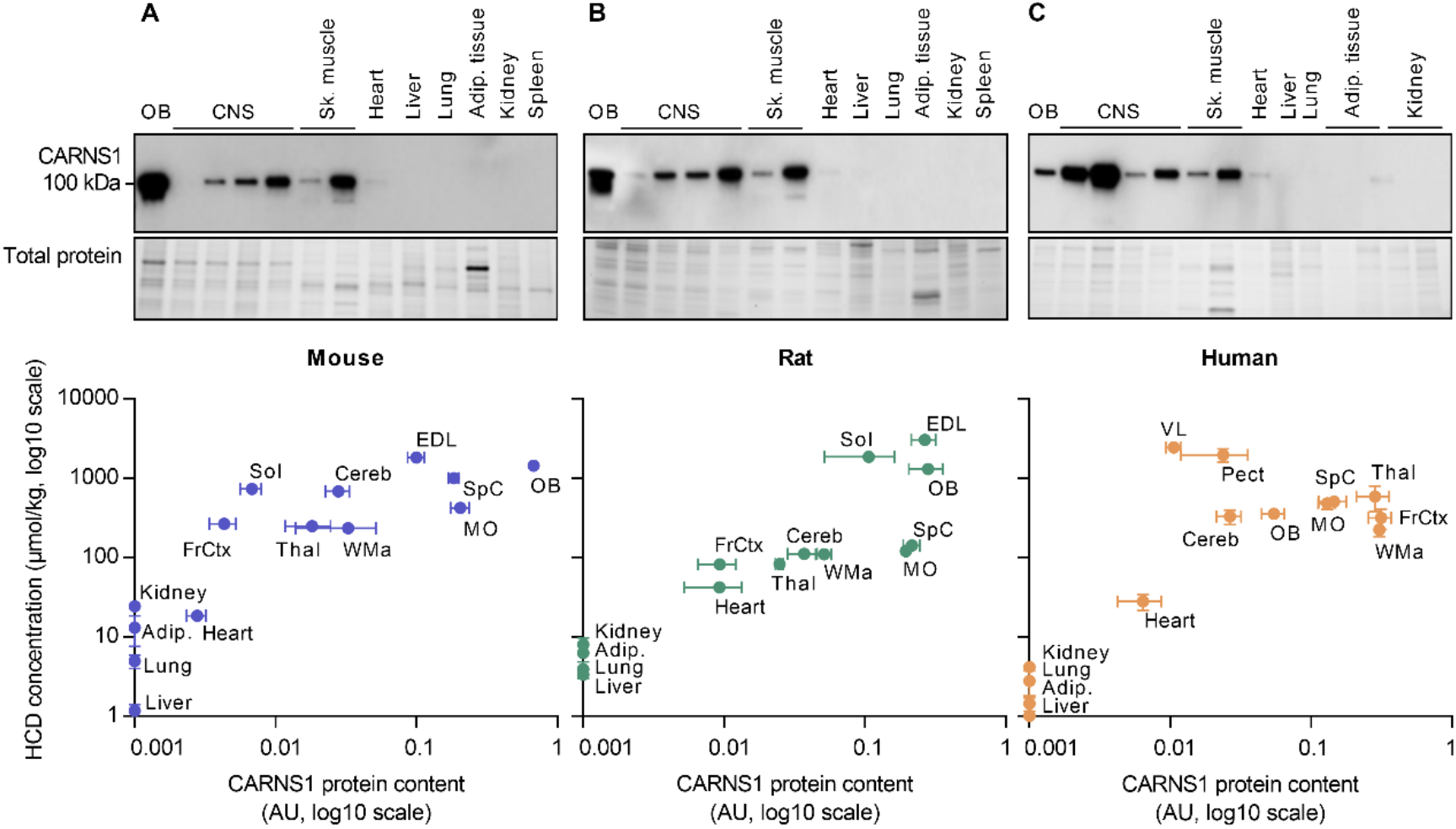
Association between CARNS1 and tissue histidine-containing dipeptide levels. Plotted relationship between CARNS1 levels (Western blot) and HCD measurements (UHPLC-MS/MS) in a variety of **(A)** mouse, **(B)** rat, and **(C)** human tissues. The central nervous system (CNS) regions that are shown, besides the olfactory bulb (OB), are frontal cortex (FrCtx), cerebellum (Cereb), spinal cord (SpC), thalamus (Thal), white matter (WMa) and medulla oblongata (MO). Mouse muscles are soleus (Sol) and extensor digitorum longus (EDL). Human muscles are m. vastus lateralis (VL) and m. pectoralis (Pect). Human adipose tissue is subcutaneous and visceral fat (Adip). Human kidney is medulla and cortex. Data are mean ± SEM. AU arbitrary units.

### N-acetylcarnosine and balenine are mainly found in human skeletal muscle

Whilst the parent HCDs (carnosine and homocarnosine) were ubiquitously expressed, most of the examined tissues also contained at least one methylated (anserine or balenine) or acetylated (N-acetylcarnosine) carnosine analog (**Fig 1F**). Rodent skeletal muscles contained by far the highest anserine levels (up to ~1500 μmol/kg in the rat EDL, **Fig 6A**). Besides anserine (~40 μmol/kg, **Fig 6A**), human skeletal muscle also contained N-acetylcarnosine (~25 μmol/kg, **Fig 6B**) and balenine (~5 μmol/kg, **Fig 6C**). In fact, human skeletal muscle was the tissue where we observed the highest N-acetylcarnosine and balenine levels. **Fig 6D-F** display the proportion of methylated or acetylated carnosine variants in different species and tissues.

**Figure 6.**
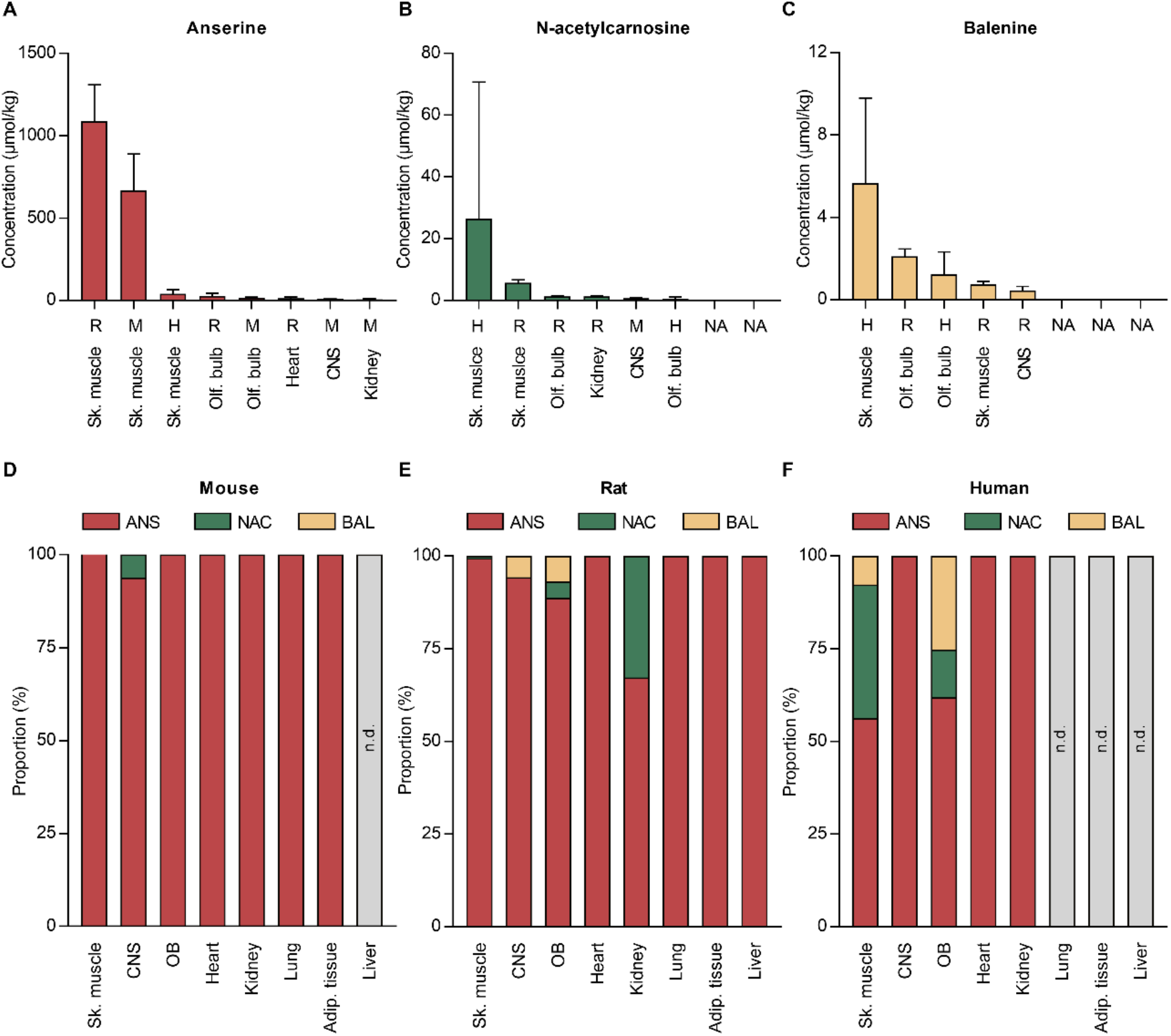
Anserine, N-acetylcarnosine and balenine levels in mouse, rat and human tissues. **(A)** Anserine, **(B)** N-acetylcarnosine, and **(C)** balenine measurements by UHPLC-MS/MS in mouse (M), rat (R), and human (H) tissues. The figures display the 8 tissues with the highest concentration of each dipeptide. Data are mean ± SD. (**D-F**) Relative proportion of anserine, N-acetylcarnosine, and balenine in **(D)** mouse, **(E)** rat, and **(F)** human tissues. Adip., adipose; ANS, anserine; BAL, balenine; CNS, central nervous system; NAC, N-acetylcarnosine; n.d., not detectable; Olf. bulb olfactory bulb; Sk., skeletal.

### Acetylation of carnosine provides stability in the human circulation, which is not required in red blood cells or rodents

Since rodents and humans differ substantially in presence and activity of the hydrolyzing enzyme CN1 in the circulation (*2, 10*), we attempted to map the circulating content of the five main HCDs. As expected, levels of plasma carnosine and anserine were very high in mice (~1500 nM) and rats (~3500 nM), but in the low nanomolar range in humans (**Fig 7A**). Low levels of homocarnosine could be detected in all 3 species, while balenine was only present in human plasma (**Fig 7A**). Interestingly, N-acetylcarnosine was the most abundant HCD in the human circulation, accounting for ~45% of the total HCDs (**Fig 7B**). Although HCD levels in red blood cells (RBCs) were in the nanomolar range in all 3 species, striking differences were observed compared to plasma (**Fig 7A** and **7C**). For both rodent species, HCD levels were drastically lower in RBCs than plasma (**Fig 7D** and **S3**). On the contrary, human carnosine and anserine levels were higher in every RBC sample compared to plasma, whilst N-acetylcarnosine levels were lower in RBCs than plasma (**Fig 7E**). These data suggest that carnosine in the human circulation is rendered more stable via acetylation to N-acetylcarnosine (which is resistant to hydrolysis by CN1) or via transport inside RBCs.

**Figure 7.**
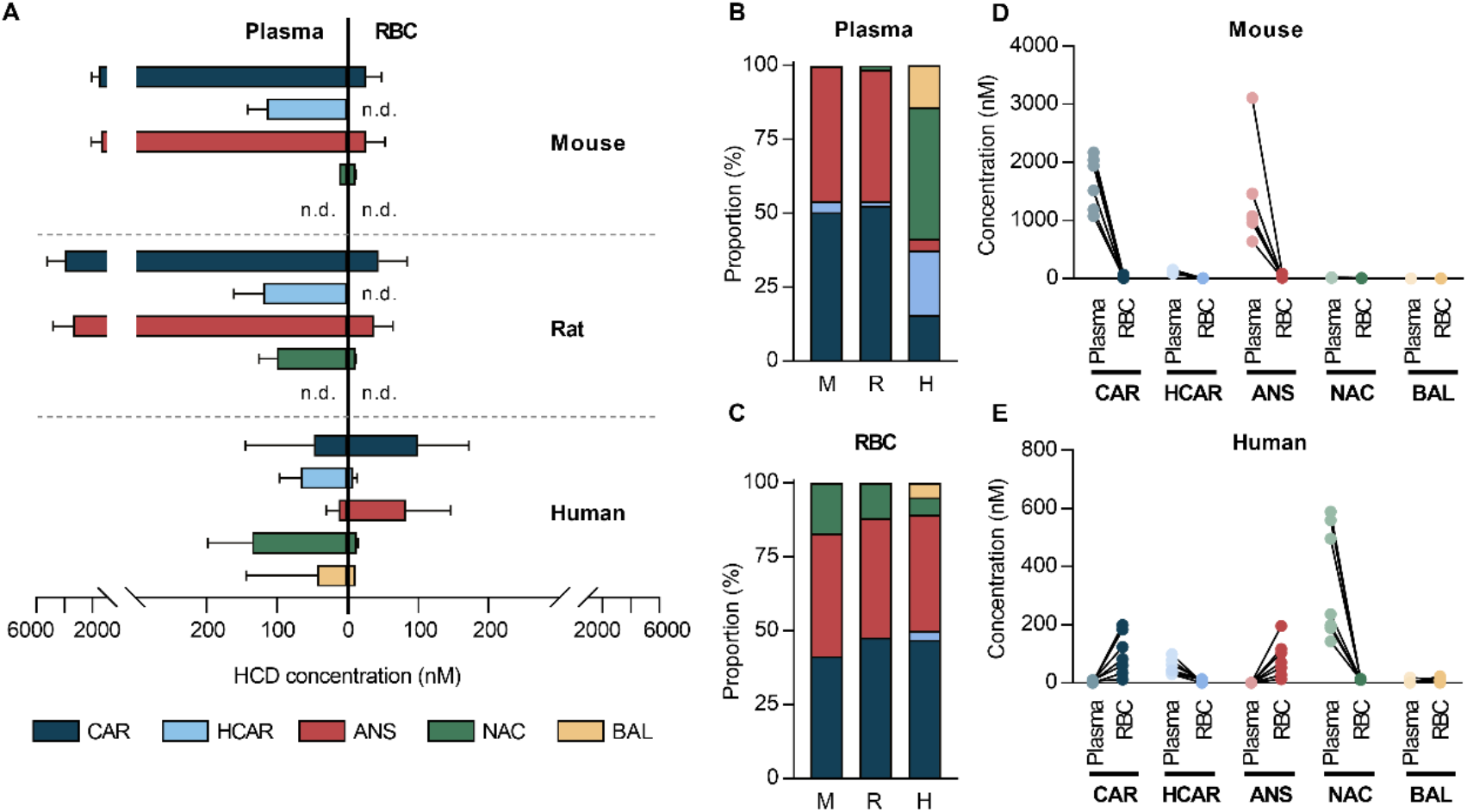
Species differences in histidine-containing dipeptides in plasma and red blood cells. **(A)** HCD measurements by UHPLC-MS/MS in mouse, rat and human plasma and red blood cells (data are mean ± SD). **(B)** Relative proportion of each HCD in mouse (M), rat (R) and human (H) plasma. **(C)** Relative proportion of each HCD in mouse, rat and human red blood cells. **(D)** Direct comparison of HCDs in plasma and red blood cells collected from the same mice. **(E)** Direct comparison of HCDs in plasma and red blood cells collected from the same human individuals. ANS anserine; BAL, balenine; CAR, carnosine; HCAR, homocarnosine; NAC, N-acetylcarnosine; RBC, red blood cells.

### Oral β-alanine supplementation affects HCDs in skeletal muscle and circulating N-acetylcarnosine levels in humans

We next investigated the effects of chronic supplementation of the rate-limiting precursor β-alanine on the content of all five HCDs in human skeletal muscle. As expected, β-alanine supplementation increased muscle carnosine content (+89%, **Fig 8A** and **S4A**). In addition, we observed increases in muscle anserine (+38%, **Fig 8A** and **S4B)**, N-acetylcarnosine (+97%, **Fig 8A** and **S4C**) and balenine (+199%, **Fig 8A** and **S4D)**, and these increases were proportional to the carnosine increase (**Fig 8B**). However, homocarnosine content significantly decreased (−50%) after 12 weeks of β-alanine supplementation (**Fig 8A** and **S4E**). Plasma N-acetylcarnosine increased by 48% after the supplementation period (**Fig 8A** and **S4F**), which correlated at the individual level with muscle carnosine content (**Fig 8C**), suggesting a possible link between intramuscular and circulating HCD levels. In summary, oral β-alanine supplementation is a potent stimulus affecting all HCDs in skeletal muscle, and plasma N-acetylcarnosine may reflect muscle HCD levels under baseline conditions and during β-alanine supplementation.

**Figure 8.**
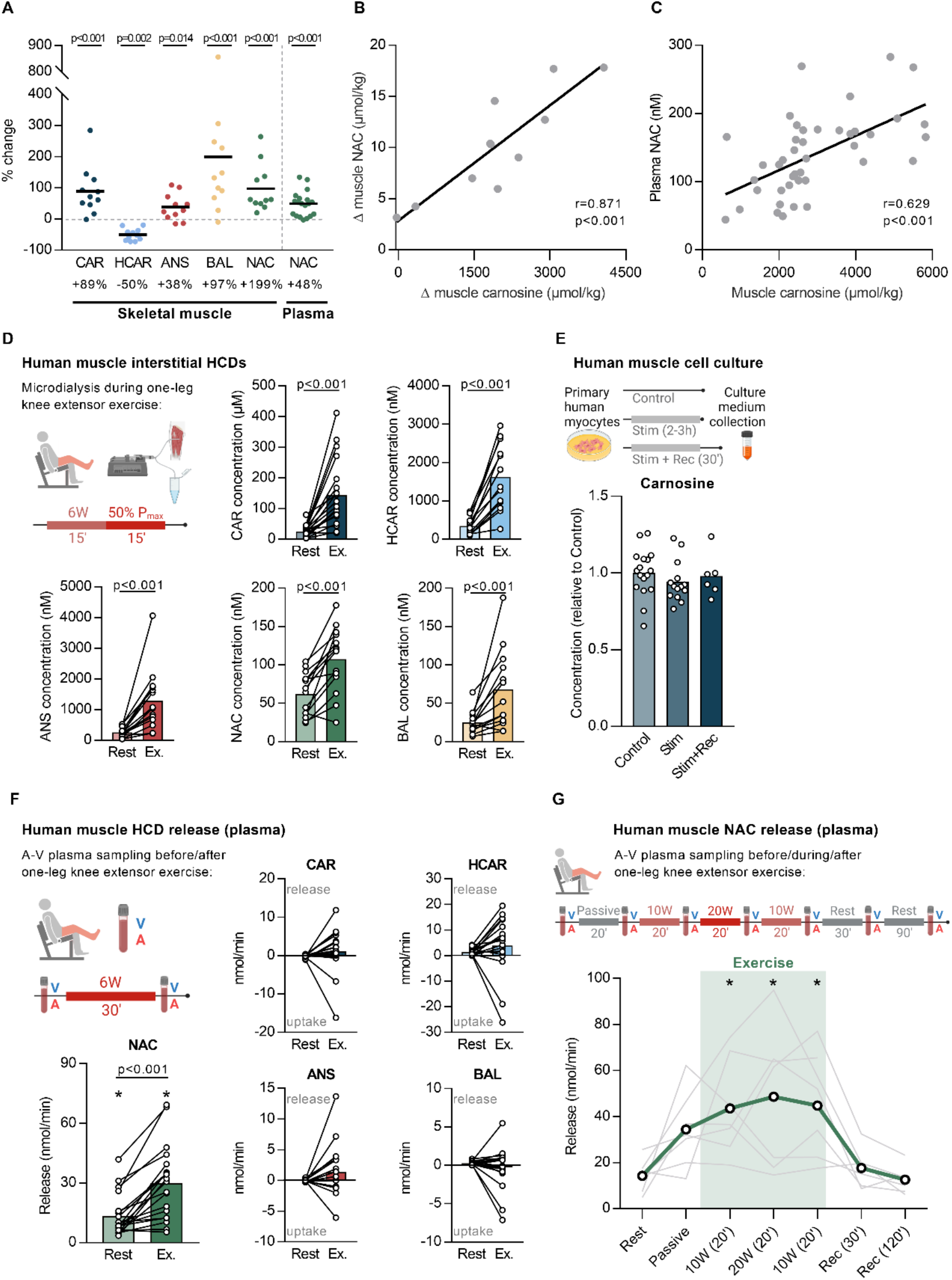
β-alanine supplementation and N-acetylcarnosine release during exercise from human muscle. **(A)** Changes in HCD levels, measured by UHPLC-MS/MS, in human skeletal muscle (m. vastus lateralis) after β-alanine supplementation. **(B)** Correlation between supplementation-induced changes in skeletal muscle carnosine and N-acetylcarnosine, measured by UHPLC-MS/MS. **(C)** Correlation between skeletal muscle carnosine and plasma N-acetylcarnosine, measured by UHPLC-MS/MS. **(D)** HCD measurements by UHPLC-MS/MS in human skeletal muscle interstitial fluid at rest and following exercise (Ex.). **(E)** Carnosine measurements by UHPLC-MS/MS in culture medium from primary human muscle cells in control condition, after 2-3 h of electrical stimulation (Stim), and after 2-3 h of electrical stimulation followed by 30 min recovery (Stim+Rec). **(F)** HCD measurements by UHPLC-MS/MS in human arterial and venous plasma samples at rest and following exercise (Ex.). Positive values indicate a net release, negative values indicate a net uptake. Asterisk indicates significantly different from zero release at the respective time point. **(G)** Release of N-acetylcarnosine in human plasma, based on arterio-venous differences, at rest, during passive movement, at different time points during exercise, and up to 120 min recovery (Rec). Positive values indicate a net release, negative values indicate a net uptake. ANS anserine; A-V; arterio-venous, BAL, balenine; CAR, carnosine; HCAR, homocarnosine; NAC, N-acetylcarnosine.

### N-acetylcarnosine is released from human skeletal muscle during exercise

Given this likely relationship between intra- and extracellular HCDs, and given that skeletal muscle is the main active organ during exercise, we explored HCD dynamics during exercise. First, we collected muscle interstitial fluid at rest and during exercise in humans. During exercise, interstitial levels for every HCD increased (**Fig 8D**). This increase could however be primarily caused by sarcolemmal damage following insertion of the microdialysis probe (*19*). To check this, we performed two follow-up experiments. Firstly, *in vitro* human primary muscle cells were electrically stimulated for 2-3 h to simulate muscle contraction. This did not result in secretion of carnosine into the culture medium immediately after the electrical stimulation or following 30 min recovery (**Fig 8E**). Other HCDs could not be detected in the cell culture medium. Secondly, we collected interstitial fluid from mouse skeletal muscle, with a previously published method in which the muscle is not mechanically affected (*20, 21*). Interstitial levels of carnosine and anserine were not higher in exercised mice compared to control mice (**Fig S5A**). Next, we collected samples of the femoral artery and vein at rest and during exercise (from a group of postmenopausal women). Our results clearly indicate a release of N-acetylcarnosine from muscle tissue at rest of ~13 nmol/min (from one leg), which further increased ~2-fold during exercise (**Fig 8F**). No release at rest or during exercise was observed for any of the other HCDs (**Fig 8F**). These results were confirmed using a similar experimental setup in healthy young men, and with more sampling time points during rest, passive movement, exercise and recovery. Here, we again showed a release of N-acetylcarnosine at rest (~14 nmol/min from one leg), which increased 3.4-fold during exercise and quickly returned back to baseline during recovery (**Fig 8G**). Exercise did not induce carnosine release or uptake within RBCs (**Fig S5B**). Also in mice, no changes in carnosine or anserine levels within plasma were observed following 60 min exercise (**Fig S5C**). Taken together, these data indicate that N-acetylcarnosine in humans is likely the major, or most stable, HCD released during exercise in a myokine-like fashion.

## Discussion

This is the first study to systematically and extensively profile the organ distribution of HCDs and their differences between mice, rats and humans. Although present in all investigated non-excitable tissues in minute amounts, mainly excitable tissues contained high HCD levels in all species. Yet, our data show surprisingly low values in cardiac tissue across species, and a different distribution of HCDs in CNS regions between species. The enzyme CARNS1 is the unique enzyme responsible for endogenous carnosine and homocarnosine synthesis, and is a major determinant for tissue HCD levels. In human CNS white matter, CARNS1 appears restricted to cells from the oligodendrocyte lineage. We also uncover that N-acetylcarnosine is the primary circulating HCD in human plasma and is continuously secreted from skeletal muscle into the circulation, which is further increased by physical exercise. An overview of the main findings is visualized in **Fig 9**.

**Figure 9.**
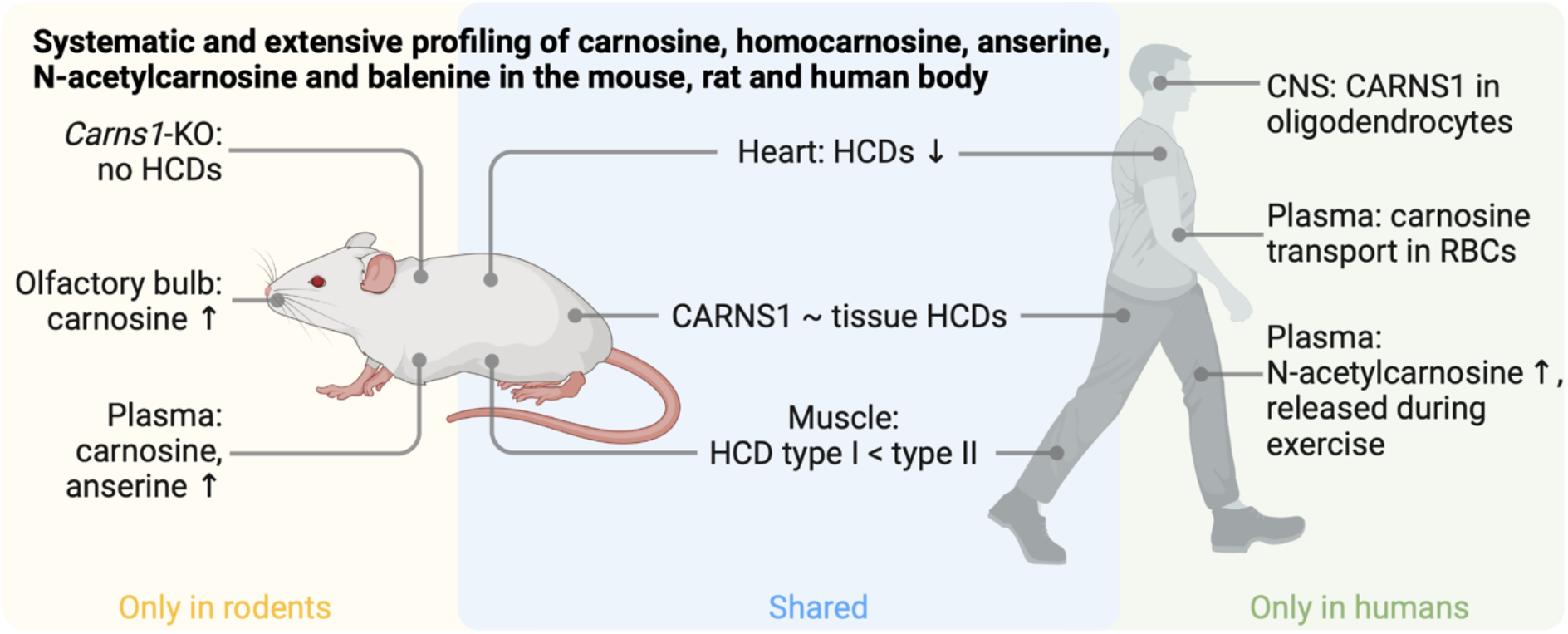
Graphical summary. Visual representation of the main findings, based on our extensive profiling of histidine-containing dipeptides (HCDs) in mice, rats and humans, and related follow-up experiments. Arrow up indicates high abundance, arrow down indicates low abundance. CNS central nervous system; RBC, red blood cell.

Previous endeavors to profile HCDs mostly focused on rodent tissues, and resulted in fragmented and sometimes contradictory literature (*13, 22–25*). Our systematic approach and very sensitive state-of-the-art UHPLC-MS/MS methodology facilitate direct comparison between tissues and species. Our data contradicted some of the previous findings, e.g. that human muscle contains only carnosine and no other HCDs (*2*), that rat kidney, lung, plasma and liver lack HCDs (*13*), or that homocarnosine is exclusively found in the CNS (*26*), with no presence of carnosine in the human brain or cerebrospinal fluid (*27*). We confirmed that anserine is predominantly found in rodent skeletal muscles (*2, 16*), but add that N-acetylcarnosine and balenine were primarily enriched in human skeletal muscles, although still lower than anserine and (homo)carnosine. From our systematic approach, we calculate that 99.1% of the total amount of HCDs in the human body is found in skeletal muscle tissue, which confirms previous estimations (*28*). Moreover, HCD concentrations in the present study are often different than those previously reported. More specifically, we report lower HCD concentrations in human skeletal muscle than previous studies (*29–31*). This is most likely explained by the use of tissue-specific *Carns1*-KO tissue matrix for our quantification method, which is known to be important for MS-based quantification (*32*). Furthermore, Peters *et al*. reported very high carnosine (1800 μmol/kg) and anserine (4000 μmol/kg) concentrations in human kidney (*23*), which are approximately 1000-fold higher than the concentrations in our dataset (~2 μmol/kg). Besides the use of a different detection technique and quantification method, it is unclear where such differences may have originated from.

CARNS1 and HCDs, especially carnosine, have long been recognised as enriched compounds within the olfactory tract of rodents (*33, 34*). Immunostaining and mass spectrometry imaging revealed the enrichment of CARNS1 and carnosine in the outer (especially glomerular) layers of the mouse olfactory bulb. This also highlights the potential of novel imaging techniques in future HCD research. In human olfactory bulbs, we found remarkably low levels of carnosine compared to homocarnosine. In addition, this is the first study to unequivocally ascribe CARNS1 expression to a specific cell type within the CNS. Our discovery of CARNS1 localisation within cells of the oligodendrocyte lineage (human white matter) confirms suggestions from recent RNA sequencing databases of the mouse and human CNS that reported *Carns1*/*CARNS1* as an oligodendrocyte-enriched gene within brain parenchyma (*35, 36*).

Traditionally, the highest HCD levels are assigned to the excitable tissues. Though this holds true with respect to skeletal muscle and CNS tissue, it was quite compelling that we found extremely low amounts of HCDs in cardiac muscle tissue. In contrast to previous suggestions that the rat heart contains ~10 mM HCDs (*37*), we report 100-fold lower levels. This was consistently found in all three species we investigated, and is also 50- to 100-fold lower compared to the concentrations in skeletal muscle. This can potentially be attributed to more accurate and sensitive quantification compared to the older technology. Nevertheless, HCDs are thought to play a crucial role in cardiac function and recovery from injury (*12, 38*). For instance, isolated cardiac myocytes from *Carns1*-transgenic hearts were protected against hypoxia reoxygenation injury (*39*), whilst *Carns1*-KO rats have impaired cardiac contractility accompanied by reduced Ca^2+^ peaks and slowed Ca^2+^ removal (*40*). This underscores the physiological importance of HCDs and indicates that even low HCD levels can contribute significantly to cell/organ function and health. It remains to be determined, however, which biochemical properties and physiological roles of these pleiotropic molecules are the most important in different tissues and under different conditions. With respect to cardiomyocytes, it has been proposed that carnosine functions as a Ca^2+^/H^+^ exchanger to shuttle calcium towards and protons away from the sarcomere site (*41, 42*).

The parent HCDs carnosine and homocarnosine share the same synthesizing enzyme CARNS1. This also underlies our observation that oral β-alanine intake leads to increased carnosine but reduced homocarnosine levels in human muscle, implying substrate inhibition between GABA and β-alanine for CARNS1. Additionally, high expression of CARNS1 can lead to high tissue content of either carnosine or homocarnosine, probably dependent of the local precursor availability (GABA *vs*. β-alanine). *Carns1*-KO mice did not produce HCDs, whereas in WT mice, rats and humans, there appears to be a relationship between CARNS1 expression and HCD content on a whole-body level. Moreover, the differences in HCD content between slow- and fast-twitch muscle fibers were paralleled by similar differences in CARNS1 expression. However, the correlation between CARNS1 expression and HCD content was not perfect, suggesting that there might be inter-organ exchange or that there is higher HCD turnover in tissues that have an important role for carnosine consumption/recycling, for example through oxidative stress and reactions with toxic metabolites in pathological conditions (*8*). Alternatively, the fact that CARNS1 shows a preference for β-alanine compared to GABA as a substrate may skew this relationship considering that some tissues contain more carnosine than homocarnosine and vice versa (*11*).

Tissues that have no or minimal CARNS1 expression likely rely on transmembrane uptake of HCDs derived from exogenous/dietary sources or from production in CARNS1-expressing organs. It has remained unclear, however, if and how HCDs are transported between tissues. The detection of HCDs in various rodent and human tissues likely illustrates that HCDs can be exchanged between organs with and without synthesizing capacity. Indeed, mice that lack the carnosine transporter PEPT2 have altered (mostly reduced) HCD levels in various organs but increased levels in skeletal muscle tissue, which is capable of synthesizing carnosine itself (*22*). This issue remains largely unclear in humans, in which high activity of the CN1 enzyme quickly degrades circulating carnosine (*2*). It has long been suggested that circulating HCDs are extremely low or absent in human plasma (*43-45*), although more recent reports already detected carnosine (*46, 47*). We now demonstrate that N-acetylcarnosine is the most stable carnosine analog in plasma, indicating that acetylation of the β-alanine residue protects against the hydrolyzing activity of CN1. Thus, N-acetylcarnosine may be the primary HCD that is exchanged between tissues in humans. Interestingly, plasma N-acetylcarnosine levels were correlated to muscle carnosine (and N-acetylcarnosine) levels, possibly indicating that plasma N-acetylcarnosine can serve as a surrogate marker for intramuscular HCD levels. Plasma N-acetylcarnosine levels increased following β-alanine intake, showing that circulating N-acetylcarnosine levels may also be a marker for muscle carnosine/HCD loading. In addition, our data indicate that transport of carnosine in RBCs is an alternative strategy to protect against CN1 in human plasma, as recently suggested (*48*). This was not true for rodents, who had lower HCD levels in RBCs than humans, despite more than 25 times higher plasma HCD levels.

It has been proposed that carnosine is released from muscles during periods of contractile activity, potentially serving as a health-promoting myokine. This hypothesis is primarily based on a study in rats, where plasma carnosine levels increased during the dark/active phase when animals were provided with a running wheel (*49*). We report that in humans, N-acetylcarnosine is the only HCD that is consistently released from muscle tissue into plasma at rest, which is further increased during periods of muscular activity (exercise). This opens various new research avenues on N-acetylcarnosine as an exercise-induced myokine (*50, 51*). Future experiments should determine its relevance for exercise training adaptations and cell/organ crosstalk (*52*). The average N-acetylcarnosine release of 14.3 nmol/min from non-contracting leg muscles at rest is striking. Extrapolation of this release to the whole body, assuming that all muscles have the same N-acetylcarnosine secretion, suggests that in theory the blood N-acetylcarnosine concentration should increase by 23.9 μM every 24 hours. Despite this continuous and substantial release of N-acetylcarnosine into the circulation, resting plasma N-acetylcarnosine levels only reach 50-350 nM in most subjects, indicating that there is a large uptake/utilization of N-acetylcarnosine in other organs, or urinary excretion. Based on the amount of N-acetylcarnosine release measured in arterio-venous samples from the leg, we also estimated that a daily turnover of 25% of the total muscle N-acetylcarnosine pool is needed to maintain stable muscle N-acetylcarnosine levels, suggesting a rather dynamic HCD homeostasis in human muscle. Specific description of these calculations and used assumptions can be found in the **Supplementary Text**. In muscle interstitial fluid samples, all HCDs appear to increase during exercise. However, it is hard to distinguish physiological exercise-induced release from sarcolemmal rupture caused by insertion of the microdialysis probes (*19*), as also supported by (I) the absence of a carnosine release during electrical stimulation of primary human muscle cells or interstitial fluid sampled from mice post-exercise, and (II) the lack of other HCD release (besides N-acetylcarnosine) in the venous effluent of contracting muscles.

Despite being the first study to systematically and extensively study the distribution of HCDs in three species, we acknowledge that the mouse and rat data cannot be fully extrapolated to all mouse and rat strains, since these can differ somewhat (*53*). We also decided to focus on the two parent HCDs (carnosine and homocarnosine) and carnosine’s methylated and acetylated analogs (anserine, balenine, N-acetylcarnosine). Other HCD conjugates do exist, but these are mostly very low abundant products from reactions with other (toxic) compounds (e.g. carnosine-propanol or 2-oxo-carnosine (*54–56*)).

In conclusion, we extensively profiled the organ distribution of the five main HCDs and discovered new physiological routes of HCD metabolism. Our results can be used to generate various new research hypotheses and highlight that findings derived from animal research on HCDs can not always be translated to humans. In particular, the apparent inter-cell and inter-tissue para- and endocrine regulation of HCDs, as well as its relevance to human health, disease and exercise performance/adaptation, deserve further investigation.

## Materials and methods

### HCD profiling - Rodent tissue collection

All mouse tissues were obtained from an in-house breeding of *Carns1*-KO and WT mice with a C57/BL6 background, kindly provided by Prof. M. Eckhardt (*14, 57*). Genotypes of the offspring from heterozygous parents were determined in toe samples using a previously published protocol (*14*). Wistar rats were supplied by Envigo (The Netherlands). Mice and rats were housed under standard room conditions (12h:12h light:dark cycle, 20-24°C, relative humidity 30-60%) and had *ad libitum* access to drinking water and food pellets. For tissue collection, female and male animals were sacrificed at an age of 6-10 w (mice) or 7-8 w (rats) old. Following overdose injection of Dolethal (200 mg/kg, i.p.), blood was collected from the right ventricle, kept in Multivette^®^ 600 K3 EDTA vials on ice, centrifuged (5 min, 3500 rpm), and plasma was stored at −80°C. Before tissue dissection, mice and rats were perfused with 0.9% NaCl solution containing heparin (25 UI/mL) via a left ventricular puncture. For determination of HCD levels by UHPLC-MS/MS and CARNS1 expression by Western blot, tissues were immediately frozen in liquid nitrogen, before being stored at −80°C. For immunohistochemistry, whole mouse brains were carefully placed on a metal plate cooled by dry ice in a foam box for several minutes, wrapped in aluminum foil, and stored at −80°C. Mouse exercise experiments were performed on a treadmill (6 m/min, speed increased every 2 min by 2 m/min until 16 m/min, total duration 60 min), and were preceded by a 1-week adaptation period (3 running sessions, gradually increasing exercise intensity and duration). Sedentary mice were placed on a stationary treadmill for 60 min. Immediately after exercise, plasma was obtained and mice were perfused as described above. To collect interstitial fluid, gastrocnemius muscles were placed on 20 μM nylon net filters (Millipore, cat# NY2004700) and centrifuged (10 min, 800 × g) (*20, 21*). All animal procedures were approved by the Ethical Committee on Animal Experiments at Hasselt University (202074B, 202127 and 202145).

### HCD profiling - Human tissue collection

All human samples were obtained after written informed consent.

#### Human vastus lateralis muscle

Muscle biopsies were collected from the m. vastus lateralis of healthy, young volunteers using the Bergström needle biopsy technique with suction, as described previously (*58*). One part of the samples was immediately snap-frozen in liquid nitrogen and stored at −80°C until UHPLC-MS/MS analysis. The other part was submerged in 1-1.5 mL of RNA*later* (Thermo Fisher Scientific), stored at 4°C for maximum 48 h and subsequently stored at −80°C for later individual fiber dissection.

#### Human heart and pectoralis muscle

Heart and pectoralis muscle samples were collected from patients undergoing open heart surgery under general anaesthesia. Heart samples consisted of the right atrial appendage, harvested at the time of venous drainage cannulation for cardiopulmonary bypass. Samples were immediately snap-frozen in liquid nitrogen and stored at −80°C. For **Fig 1F**, HCD concentrations of vastus lateralis and pectoralis muscle were averaged.

#### Human kidney

Kidney samples were collected from patients undergoing radical nephrectomy. In case of kidney cancer, tissue was sampled as far away from the site of the tumor to obtain the healthiest part of the kidney, immediately snap-frozen in liquid nitrogen and stored at −80°C. Medulla and cortex were sampled separately (**Table S1**), but data was later averaged for analysis and visualization.

#### Human adipose tissue

Visceral and subcutaneous adipose tissue were sampled from lean and obese individuals during abdominal surgery following an overnight fast (*59*). Tissue pieces were rinsed, freed from visible blood and connective tissue, and snap-frozen in liquid nitrogen. The two subtypes (**Table S1**) were later averaged for analysis and visualization.

#### Human lung

Lung tissue was obtained from unused healthy donor lungs that were not used for transplantation from the BREATH KULeuven biobank (S51577).

#### Human liver

Liver samples were surgically removed from fasted subjects with obesity during gastric bypass surgery. Exclusion criteria were the presence of malignancies, drinking more than two units (women) or three units (men) of alcohol per day, having known liver pathologies other than non-alcoholic fatty liver disease. For the current analysis, five samples with NAS score 0 or 1 were selected (*60*).

#### Human plasma and RBC

Blood samples were always collected in pre-cooled EDTA tubes, after a 1-2-day lacto-ovo vegetarian diet to ensure no influence of dietary HCD intake. For plasma, samples were immediately centrifuged (10 min, 3000 × g, 4°C), followed by immediate deproteinization (110 μL of 35% 5-sulfosalisylic acid per 1 mL of plasma) and second centrifugation (5 min, 15000 × g, 4°C). Plasma samples were then stored at −80°C. For RBC isolation (*61*), blood tubes were centrifuged for 15 min at 120 × g at room temperature. Plasma was carefully removed and 200 μL RBCs were collected in 1.8 mL of ice-cold methanol (55% v/v). Samples were then stored at −80°C.

#### Human CNS

Seven different human CNS regions were obtained from The Netherlands Brain Bank (NBB), Netherlands Institute for Neuroscience, Amsterdam (open access www.brainbank.nl). All donors were ‘non-demented controls’, indicating the absence of neurological and psychiatric disease. From 2 subjects, all 7 regions were available. For immunohistochemical analyses, white matter tissue from healthy controls and unaffected white matter tissue of multiple sclerosis patients from a previous study was used.

#### Human cerebrospinal fluid

Cerebrospinal fluid from subjects without neurological disease at the time of sampling was obtained via the University Biobank Limburg (UbiLim), with approval from the Medical Ethics Committee at Hasselt University (CME2021-004). Lumbar cerebrospinal fluid was collected into PPS tubes, kept at 4°C, centrifuged to remove cells (10 min, 500 × g, 4°C), and supernatant was stored at −80°C.

### Human β-alanine supplementation study

Vastus lateralis muscle biopsies obtained via the Bergström needle biopsy technique with suction (n=11 β-alanine, n=11 placebo) and plasma samples (n=19 β-alanine, n=17 placebo) were collected before and after 12 weeks of β-alanine supplementation in patients with COPD (sustained-release CarnoSyn®, NAI). Methodological details have been described previously (*62*). Snap-frozen biopsies were freeze-dried for 48 h, followed by manual removal of non-muscle material (fat, connective tissue, blood) under a light microscope. Effects of β-alanine were analyzed using a two-way repeated measures ANOVA (group *vs*. time) for each HCD separately, followed by Sidak’s multiple comparisons tests. Correlations were performed using Pearson correlation (Δ muscle carnosine *vs*. Δ N-acetylcarnosine) or Spearman rank correlation (muscle carnosine *vs*. plasma N-acetylcarnosine).

### Human HCD release experiments

#### Microdialysis experiment

Detailed methodology has been described previously (*63*). In short, interstitial samples from m. vastus lateralis were collected using the microdialysis technique at rest and after 30 min of one-legged knee extensor exercise (15 min 6 W, 15 min 50% peak power). Subjects consisted of a group of young (n=7) and old (n=13) healthy men (results are pooled together since no differences between groups could be observed). Concentrations were corrected for probe recovery as determined by relative loss of [2–3H]-labeled adenosine in the dialysate. Interstitial levels during exercise were compared to resting values using a multiple Wilcoxon matched-pairs signed rank test with Holm-Sidak multiple comparison test.

#### Arterio-venous balance experiment 1

Arterial and venous samples were collected from the femoral artery/vein at rest and after 30min one-legged knee extensor exercise (6W) in a group of healthy postmenopausal women (n=19). Methodological details can be found in (*64*). Samples were deproteinized with 35% 5-sulfosalisylic acid, as described above, on the day of the UHPLC-MS/MS analysis. Exercise *vs*. resting HCD levels were compared using a multiple Wilcoxon matched-pairs signed rank test with Holm-Sidak multiple comparison test.

#### Arterio-venous balance experiment 2

Seven healthy, young men (28 ± 4 years old, BMI of 24 ± 2, VO_2max_ of 49 ± 7 mL/min/kg) participated in this experiment. After passive transport to the lab, catheters were inserted in the femoral artery and vein. Next, arterial and venous samples were collected every 30 min during a 90 min supine resting period. After this, 20 min of passive leg movement was performed, followed by 60 min of active one-legged knee extensor exercise (20 min 10 W, 20 min 20 W, 20 min 10 W). Samples were collected at the end of each exercise bout. Finally, arterio-venous samples were collected after 30 min and 2 h of recovery. Plasma and RBC samples were collected as described above. Exercise-induced effects were analyzed using a mixed-effect model with repeated measurements over time, with post-hoc comparison of every time point *vs*. baseline (Holm-Sidak test).

### Primary cell culture experiments

Biopsy samples (~150 mg) were obtained from m. vastus lateralis from young, healthy men. Primary skeletal muscle cells were isolated with homemade antibody-coated magnetic beads and cultured as previously described (*65, 66*). Cultured skeletal muscle cells were used for analysis on day 5 or 6 after the onset of differentiation. At this time, most of the myocytes have differentiated into multinucleated myotubes and can easily be identified as muscle cells. Myotubes were starved with media containing 0.1% Bovine Serum Albumin (DMEM without phenol, D-glucose and L-glutamine) for 16 hours before experiments. The skeletal muscle cells were electro-stimulated as described previously (*67*), with the minor addition of 5μM (S)-nitro-Blebbistatin (Cayman Chemical, CAS. 856925-75-2) to the stimulation buffer to inhibit the spontaneous contraction of the myotubes (*68*). The cells were stimulated for 2-3 h (50 Hz, 0.6s/0.4 s trains, 1 ms pulse width, 10 V, homemade electrical stimulator connected to a Digitimer MultiStim SYSTEM-D330). The extracellular medium was collected immediately or 30 min after the end of stimulation, and medium from non-stimulated control cells was harvested simultaneously. Changes over time were analyzed using a mixed-effect model with repeated measurements over time, with post-hoc comparison of stimulated and stimulated+recovery *vs*. control cells (Holm-Sidak test).

### HCD determination by UHPLC-MS/MS

Details of the validation of the in-house developed UHPLC-MS/MS analysis has been described previously (*69*), with the exception that all experiments were performed on a Xevo^®^ TQ-S MS/MS system with 2.5 μL injection volume. The limit of detection was determined to be 5-10 nM (in plasma), corresponding to 0.38-0.76 μmol/kg tissue for our homogenization protocol. Pure carnosine and anserine were kindly provided by Flamma S.p.a. (Chignolo d’Isola, Bergamo, Italy), and pure balenine by NNB Nutrition (Frisco, Texas, USA). Homocarnosine (#33695) and N-acetyl-L-carnosine (#18817) were bought from Cayman Chemical (Ann Arbor, Michigan, USA). UHPLC-MS/MS data extraction and analysis was performed using Masslynx software 4.2 (Waters, Milford, USA).

#### Tissues

All tissues were prepared similarly, based on the method described previously (*8*). Frozen tissues were quickly weighed and immediately homogenized in extraction solution (ultrapure water with 10 mM HCl and internal standard carnosine-d4) in a ratio of 95 μL extraction solution per 5 mg tissue in a QIAGEN TissueLyser II (1 min, 30 Hz). The concentration of carnosine-d4 varied according to expected HCD concentrations in the tissues: muscle (20 μM), CNS (5 μM), liver/lung/spleen (0.5 μM) and all other tissues (1 μM). Then, homogenates were centrifuged (20 min, 3000 × g, 4°C). Supernatants were immediately diluted in a 3:1 ratio with ice-cold acetonitrile (−20°C), vortexed and kept on ice for 15min. After a second centrifugation step (20 min, 3000 × g, 4°C), samples were stored at −80°C until the day of the UHPLC-MS/MS analysis. Samples were combined with 75:25 acetonitrile:water in a 4:1 ratio before injection in the UHPLC-MS/MS device. For skeletal muscle and CNS, samples were also injected after an extra initial dilution (1:25 for muscle, 1:10 for CNS) for the determination of carnosine (skeletal muscle) and homocarnosine (CNS) content. Standard calibration curves were prepared for each individual run and for each tissue separately in the respective tissue of the *Carns1*-KO mice to account for possible tissue matrix effects (for all three species). Differences between soleus and EDL HCD levels in mice and rats were analyzed using paired t-tests or Wilcoxon signed-rank tests, depending on normality of the data.

#### Plasma

Deproteinized plasma (150 μL) was combined with acetonitrile containing 1% formic acid (215 μL), 1 μM carnosine-d4 as internal standard (10 μL) and ultrapure water (25 μL). For mouse plasma analysis, volumes were scaled down to available plasma (75 μL). After thoroughly vortexing, samples were centrifuged (15 min, 15000 × g, 4°C). The supernatant was collected and injected in the UHPLC-MS/MS. A standard calibration curve was prepared in a pool of human deproteinized plasma that was collected after a 2-day lacto-ovo vegetarian diet to minimize circulating HCDs (for all three species).

#### RBC

First, 190 μL of the RBC samples was combined with 10 μL of internal standard carnosine-d4 (2μM) and 10 μL of 75:25 acetonitrile:water (with 1% formic acid). This mixture was vortexed and then centrifuged in a 10 kDa filter (Nanosep^®^ Centrifugal Device with Omega Membrane™, Pall Corporation). The supernatant was subsequently evaporated at 40°C and the droplet resuspended in 40 μL of 75:25 acetonitrile:water (with 1% formic acid) before injection in the UHPLC-MS/MS device. A standard calibration curve was prepared in a pool of human RBC samples that were collected after a 2-day lacto-ovo vegetarian diet to minimize circulating HCDs (for all three species).

#### Cerebrospinal fluid

Human cerebrospinal fluid was treated identically as plasma, but the standard calibration curve was prepared in ultrapure water since no *Carns1*-KO tissue matrix was available, possibly resulting in overestimation of the absolute concentrations.

#### Interstitial fluid

Interstitial fluid (15 μL) was combined with 30 μL acetonitrile containing 1% formic acid and 1 μM carnosine-d4 and 10 μL ultrapure water. This mixture was vortexed, centrifuged (15 min, 15000 × g, 4°C) and the supernatant was used to inject in the UHPLC-MS/MS. For detection of carnosine in humans, samples were first diluted 1:30. For detection of carnosine and anserine in mice, samples were first diluted 1:500. A standard calibration curve was prepared in Ringer-Acetate buffer as this was used to perfuse the microdialysis probes.

#### Cell culture medium

Extracellular medium (150 μL) was mixed with acetonitrile containing 1% formic acid (240 μL) and internal standard carnosine-d4 (2 μM, 10 μL), vortexed and injected in the UHPLC-MS/MS. A standard calibration curve was prepared in DMEM culture medium.

### CARNS1 protein levels by Western blot

Tissues were diluted in RIPA buffer (300 μL per 10 mg tissue; 50 mM Tris pH 8.0, 150 mM NaCl, 0.5% sodium deoxycholate, 0.1% SDS, 1% Triton-X100, and freshly added protease/phosphatase inhibitors [Roche]), and homogenized using stainless steel beads and a QIAGEN TissueLyser II (shaking 1 min, 30 Hz). Following centrifugation (15 min, 12000 × g, 4°C), supernatants were stored at −80°C. Pierce™ BCA Protein Assay Kit (Thermo Fisher) was used according to manufacturer’s instructions to determine protein concentrations (read at 570 nm wavelength). For detection of CARNS1 protein levels, 1 μg of protein was diluted in loading buffer solution (63 mM Tris Base pH 6.8, 2% SDS, 10% glycerol, 0.004% Bromophenol Blue, 0.1 M DTT), heated for 4 min at 95°C, and separated in polyacrylamide gels (4-15%, Mini-PROTEAN TGX, Bio-Rad) at 100-140 V on ice. Next, stain-free gels were imaged following UV exposure to visualise total protein content (ChemiDoc MP Imaging System, Bio-Rad). Proteins were transferred from the gel to an ethanol-immersed PVDF membrane in transfer buffer (30 min, 25 V, 1.0 A, Trans-Blot Turbo Transfer System, Bio-Rad). Membranes were briefly washed in Tris-buffered saline with 0.1% Tween20 (TBS-T), and blocked for 30 min using 3% milk powder in TBS-T. Following overnight incubation at 4°C with primary antibodies against CARNS1 (rabbit polyclonal, 1:1000 in 3% milk/TBS-T, HPA038569, Sigma), membranes were washed (3 × 5 min), incubated with secondary HRP-conjugated goat anti-rabbit antibodies (1:5000 in 3% milk/TBS-T) for 60 min at room temperature, washed again (3 × 5 min), and chemiluminescent images were developed in a ChemiDoc MP Imaging System (Bio-Rad) using Clarity Western ECL substrate (Bio-Rad). CARNS1 protein bands were quantified with Image Lab 6.1 software (Bio-Rad), and normalized to total protein content from the stain-free image. Finally, bands were expressed relative to total CARNS1 expression (sum of all bands) of a particular mouse, rat, or human. For muscle, band densities were expressed as fold changes relative to the average density from soleus muscles per blot. Linearity of the signal was determined for every tissue. Differences between soleus and extensor digitorum longus CARNS1 content were analyzed using a paired t-test (rat) or Wilcoxon signed-rank test (mouse), depending on normality of the data.

### CARNS1 protein level in human single muscle fibers

CARNS1 protein levels in type I *vs*. type II muscle fibers were determined based on a previously published method (*70*). Muscle samples in RNA*later* (Thermo Fisher Scientific) were thawed and subsequently transferred to a petri dish filled with fresh RNA*later* (Thermo Fisher Scientific) solution. Individual muscle fibers were manually dissected under a light microscope and immediately submerged in a new 0.5 mL tube with 5 μL ice-cold Laemmli buffer (125 mM Tris-HCl (pH 6.8), 10% glycerol, 125 mM SDS, 200 mM DTT, 0.004% bromophenol blue). Tubes were then incubated for 15 min at 4°C, followed by 10 min at 70°C and then stored at −80°C until the next step of the analysis. A total of 40-72 fibers were isolated from 9 biopsies. Next, muscle fiber type (based on myosin heavy chain expression) was determined using dot blotting techniques. For this, 0.5 μL of the muscle fiber lysate was spotted onto two activated and equilibrated PVDF membranes, one for MHC I and one for MHC IIa. After air-drying the PVDF membrane for 30 min, it was re-activated in 96% ethanol and equilibrated in transfer buffer (8 mM Tris-base, 39 mM glycine, 0.015% SDS, 20% ethanol). The next steps are similar to standard Western blotting as described above, with primary antibodies for MHC I (A4.840, 1:1000 in 3% milk in TBS-T, DSHB) or MHC IIa (A4.74, 1:1000 in 3% milk in TBS-T, DSHB). Fiber lysates were classified as type I or IIa fibers based on a positive stain for only the MHC I or IIa antibody (**Fig S1**). Next, fibers of the same type from the same subject were pooled (n=10-26 for type I and n=9-22 for type IIa fibers). CARNS1 protein content in these fiber type-specific samples were determined with standard Western blotting technique as described above, with loading of 5 μL per pool. Band densities were expressed as fold changes relative to the average density from type I fibers per blot. Statistical analysis was performed using a Wilcoxon signed-rank test.

### *CARNS1* mRNA expression

The fiber type-specific RNAseq dataset of Rubenstein *et al*. was downloaded from the GEO repository under accession number GSE130977 (*18*). This dataset consists of RNAseq data of pools of type I or type II fibers from m. vastus lateralis biopsies of 9 healthy, older men. For full details on generation of this dataset, we refer to the original publication. Raw counts were normalized with DESeq2 to allow for between-sample comparisons (*71*). First, normalized counts for *MYH2* (ENSG00000125414, type II fibers) and *MYH7* (ENSG00000092054, type I fibers) were extracted and assessed for each fiber pool as purity quality control. Based on this analysis, the fiber pools of one participant were excluded for further analysis. Next, normalized counts of *CARNS1* (ENSG00000172508) were extracted and compared between type I and type II fiber pools within each participant using a Wilcoxon signed-rank test.

### Immunohistochemical detection of CARNS1

Sagittal and frontal cryosections (10 μm) were cut from whole mouse brains and human white matter samples. Following acetone fixation (10 min) and blocking (30 min, 10% donkey serum in PBS with 1% BSA), sections were exposed overnight at 4°C to antibodies detecting CARNS1 (rabbit polyclonal, 1:100 in PBS with 1% BSA, HPA038569, Sigma). The next day, sections were washed and exposed to complementary secondary antibodies for 60 min (1:500 in PBS with 1% BSA, Thermo Fisher). Fluorescence imaging was performed with a Leica DM4000 B LED (Leica Microsystems). For double-labeling, we used the following antibodies: OLIG2 (1:50, goat polyclonal, AF2418, R&D Systems), neurofilament heavy polypeptide (NF-H, rabbit polyclonal, 1:200, ab8135, Abcam), CD68 (mouse monoclonal, 1:100, M0814 KP1 clone, Dako), GFAP (mouse monoclonal, 1:100, G3893, Sigma). Absence of CARNS1 from *Carns1*-KO brain sections was used as negative control.

### Matrix-assisted laser desorption ionization mass spectrometry imaging (MALDI-MSI)

The spatial distribution of carnosine and homocarnosine was studied in mouse brains by MALDI-MSI. Fresh DHB solution (20 mg/mL in 70% MeOH with 0.2% Trifluoroacetic acid) was sprayed using TM-sprayer (HTX Technologies) in a series of 15 layers with the settings: temperature 75°C, pressure 10 psi, flow rate 0.12 mL/min, velocity 1200 mm/min, track spacing 2 mm. MS acquisition was conducted on a Tims-TOF mass spectrometer (Bruker Daltonik GmbH). The data were acquired at a raster size of 50 × 50 μm (or 20 × 20 μm for higher spatial resolution) in the mass range of 100-800 *m/z* in positive ion mode (300 laser shots per pixel with 5000 Hz frequency). After performing the MALDI-MSI experiments, the matrix was gently removed by submersion in EtOH for 2 min. Slides were then washed in serial baths containing 100% EtOH, 90% EtOH, 70% EtOH or ultrapure water for 3 min each. The sections were stained by hematoxylin for 3 min, and eosin for 20 seconds. Following dehydrating steps, digital images were acquired with the Mirax system (Carl Zeiss) at 40× magnification and uploaded on Aperio ImageScope (Leica Biosystems).

### Statistical analysis

Statistical analyses were performed in GraphPad Prism v9.4 and were described in the appropriate method paragraphs. Significance level was set at α=0.05.

## Acknowledgements

For human brain tissues, all material has been collected from donors for or from whom a written informed consent for a brain autopsy and the use of the material and clinical information for research purposes had been obtained by the Netherlands Brain Bank (NBB). The experiment protocols and methods used for analysing brain samples were conducted with the approval of the NBB and the Medical Ethical Committee of Hasselt University, and carried out according to institutional guidelines. All HCDs analyses were performed using an UHPLC-MS/MS instrument part of the Ghent University MSsmall Expertise Centre for advanced mass spectrometry analysis of small organic molecules. Also the University Biobank Limburg (UBiLim) is acknowledged for providing storage and release of some human biological material used in this publication. Figure 7D, 7E, 7F and FG (experimental setups) were created in BioRender. The technical assistance of Jens Jung Nielsen, Anneke Volkaert, Thomas Ehlers, Nicklas Frisch and Josephine Rol is greatly appreciated.

## Author contributions

TVDS, JSp, SDJ and WD designed the study.

TVDS, JSp, SDJ, CH, JSt, BVer, CVA and MV performed the experiments and/or biochemical analyses.

TVDS JSp, SDJ, JDB, RVT, KV, DH, TB, BL, CVP, KD, BVan and LG contributed to tissue collection.

SC, BCP, BO, LG, YH and WD supervised the study.

TVDS JSp, SDJ and WD analyzed the data.

TVDS, JSp and WD drafted the manuscript.

All authors approved the final version of the manuscript.

## Data and materials availability

All data needed to evaluate the conclusions in the paper are present in the manuscript and supplementary materials. Additional data related to this paper can be obtained upon reasonable request to the author.

