## Supplementary Materials for "Extensive profiling of histidine-containing dipeptides reveals species- and tissue-specific distribution and metabolism in mice, rats and humans"

**This PDF file includes:**

Supplementary Text

Figs. S1 to S5

Table S1

N-acetylcarnosine release calculations

Calculations on HCD release data were based on the following assumptions:

(1) an average person of 75 kg

(2) a similar release from all muscles in the human body

(3) The whole leg is supplied with blood by a. femoralis

(4) an average release of N-acetylcarnosine from human muscle of 14.3 nmol/min from one leg (**Fig 8G**)

(5) an average concentration of N-acetylcarnosine in human vastus lateralis of 16.4 µmol/kg (**Table S1**)

(6) 40% of total body weight is muscle, and average person has 5 L of blood

(7) Fat-free mass of legs is 34.5% of whole-body fat-free mass (own unpublished data)

(8) Muscle mass of upper leg is 76% of total leg muscle mass (own unpublished data)

(9) Muscle mass of knee-extensors is 36% of total leg muscle mass (own unpublished data)

Total N-acetylcarnosine release per day from 1 leg (since a. femoralis supplies whole leg) is 20.6 µmol, or 41.2 µmol from both legs. Extrapolation to whole-body level yields 119.4 µmol, resulting in a blood concentration of 23.9 µM.

One leg of an average person weighs 5.2 kg, containing 82 µmol. A release of 20.6 µmol per day from one leg, indicates that 25.1% of the total muscle N-acetylcarnosine pool is released into the circulation per day.

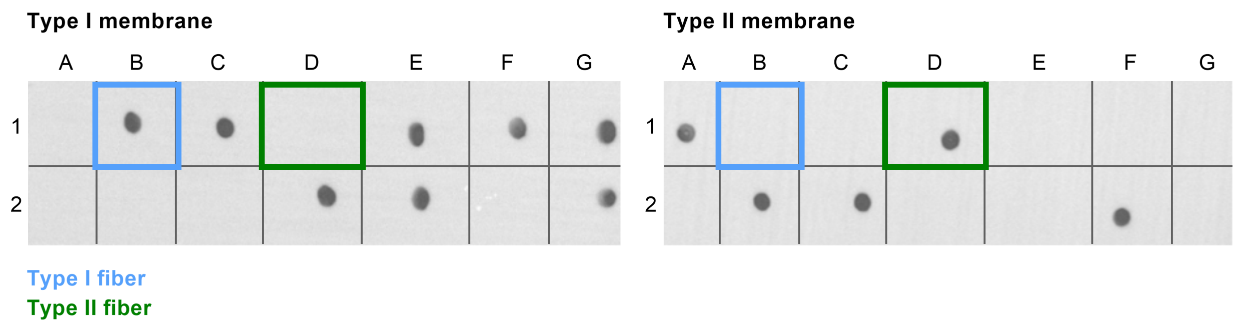


**Figure S1. Dot blot to determine muscle fiber type.** Representative image showing the distinction between type I and type II human single muscle fibers.

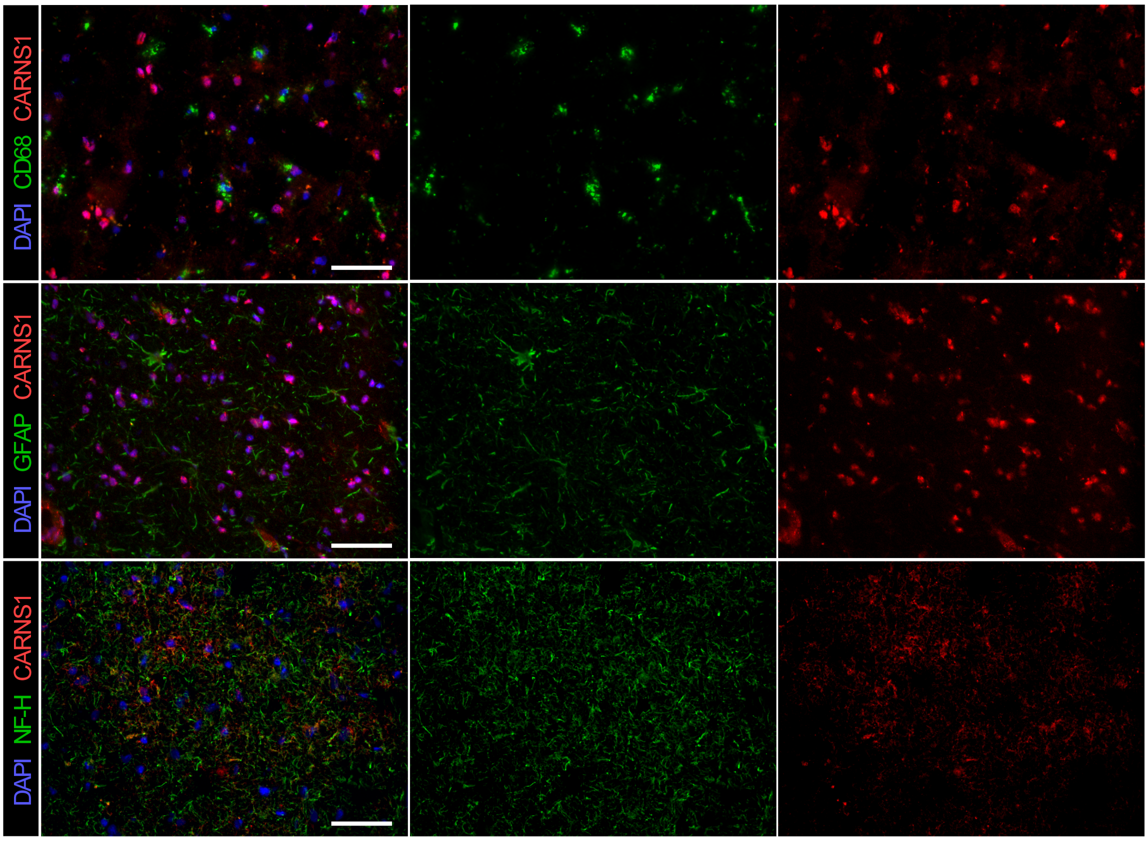


**Figure S2. CARNS1 in the human central nervous system.** Immunohistochemical detection of CARNS1 and CD68, glial fibrillary acidic protein (GFAP), or neurofilament heavy polypeptide (NF-H) in human white matter. Scale bars are 50 µm.

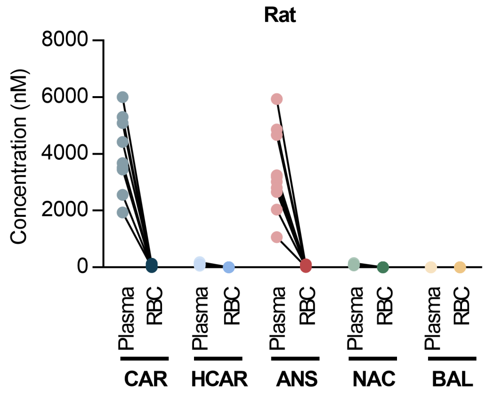


**Figure S3. Histidine-containing dipeptides in plasma and red blood cells from rats.** HCD measurements by UHPLC -MS/MS in rat plasma and red blood cells. Direct comparison of HCDs in plasma and red blood cells collectedfrom the same rats. ANS, anserine; BAL, balenine; CAR, carnosine; HCAR, homocarnosine; NAC, N-acetylcarnosine; RBC, red blood cells.

-

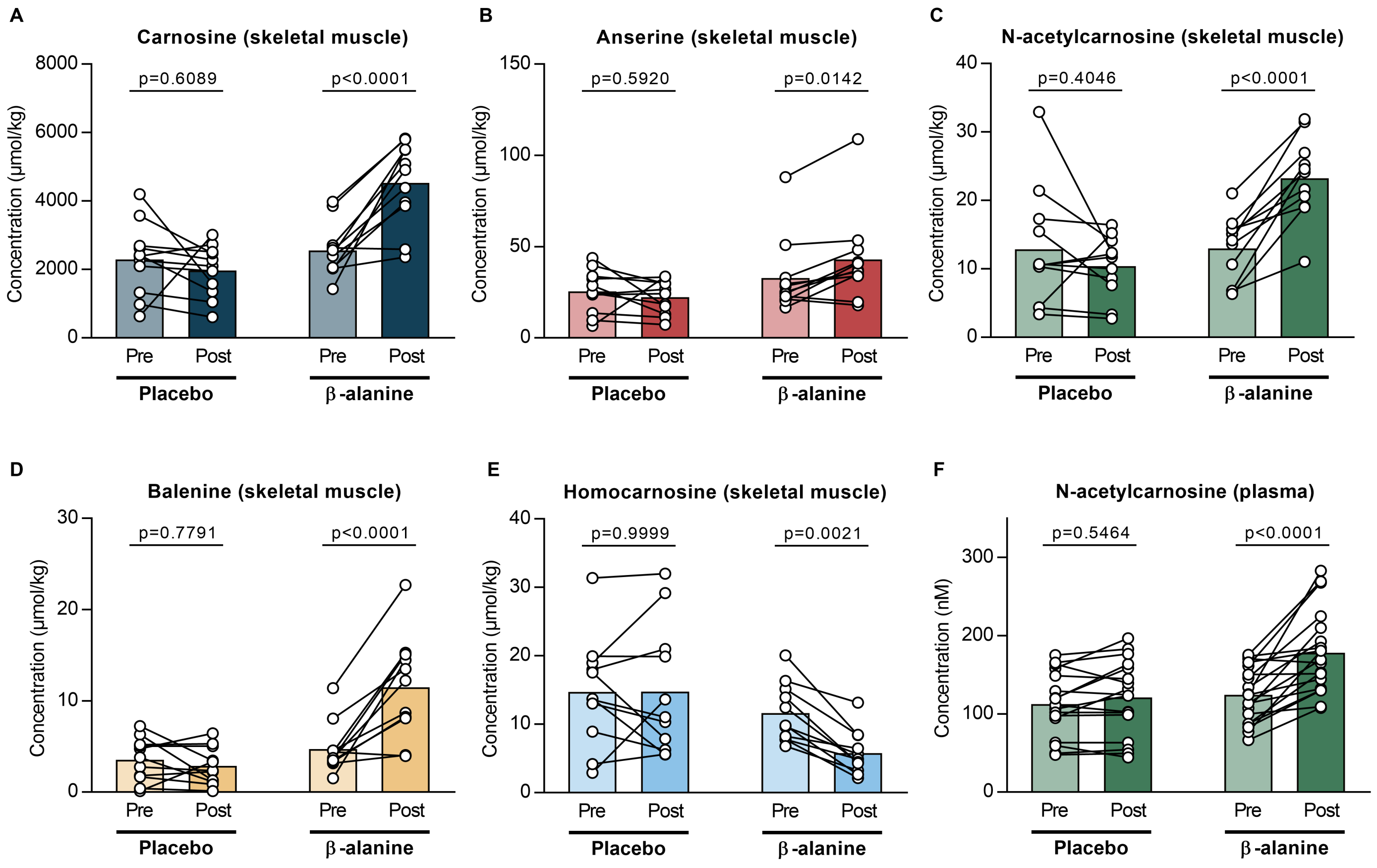


**Figure S4. Histidine-containing dipeptides in human skeletal muscle tissue and plasma following β-alanine supplementation.** UHPLC-MS/MS measurements of **(A)** carnosine, **(B)** anserine, **(C)** N-acetylcarnosine, **(D)** balenine and **(E)** homocarnosine in human skeletal muscle (m. vastus lateralis) and **(F)** N-acetylcarnosine in plasma before and after 12 weeks of β-alanine supplementation.

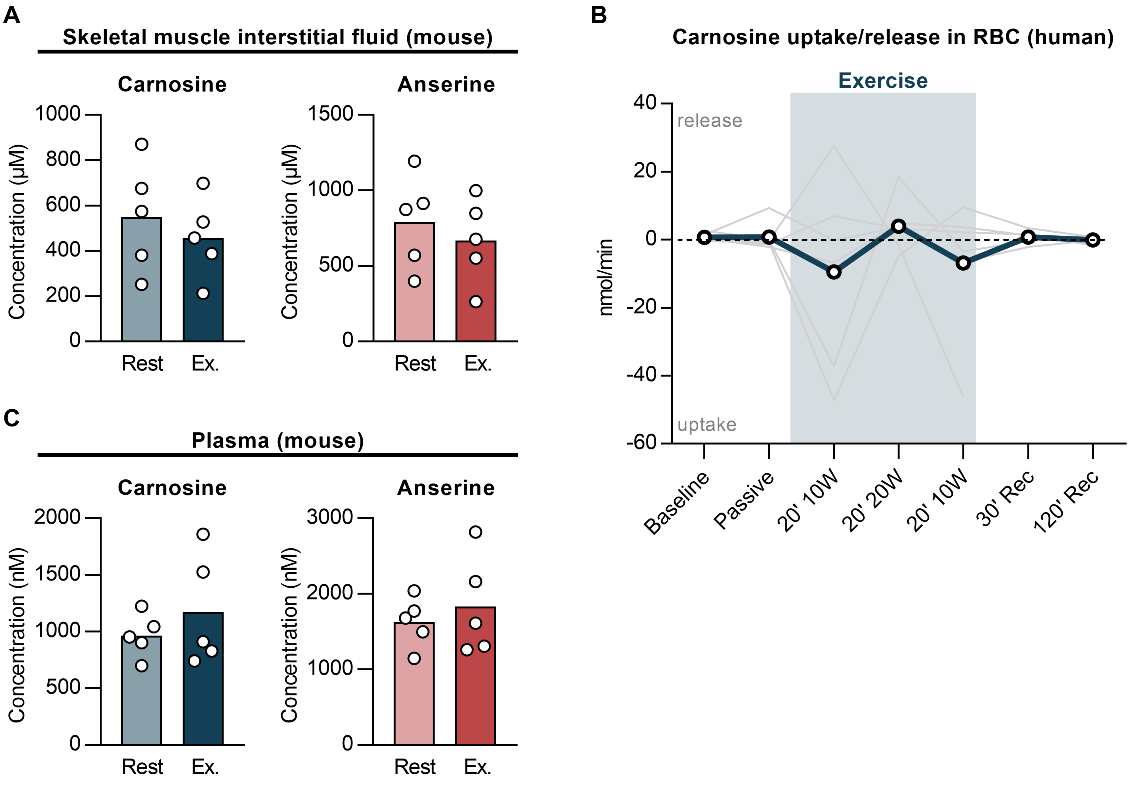


**Figure S5. HCD release from muscle during exercise. (A)** Carnosine and anserine measurements by UHPLC -MS/MS in mouse skeletal muscle interstitial fluid samples at rest and following exercise (Ex.). **(B)** Uptake or release of carnosine in human red blood cells, based on arterio-venous differences, at rest, during passive movement, at different time points during exercise, and up to 120 min recovery (Rec). Positive values indicate a net release, negative values indicate a net uptake. **(C)** Carnosine and anserine measurements by UHPLC -MS/MS in mouse plasma samples at rest and following exercise (Ex.). RBC, red blood cells.

**Table S1. Overview of histidine-containing dipeptides in the mouse, rat and human body.**

| Skeletal & cardiac muscle | | Mouse | Rat | Human |
|---|---|---|---|---|
| Soleus | Carnosine | 250 ± 168 (n=11) | 1172 ± 208 (n=10) | NA |
| | Homocarnosine | 11.9 ± 6.15 (n=11) | 17.4 ± 2.35 (n=10) | |
| | Anserine | 468 ± 195 (n=11) | 656 ± 99.8 (n=10) | |
| | Balenine | n.d. | 0.58 ± 0.23 (n=9/10) | |
| | N-acetylcarnosine | n.d. | 9.52 ± 1.84 (n=10) | |
| Extensor Digitorum Longus | Carnosine | 918 ± 287 (n=11) | 1456 ± 183 (n=10) | NA |
| | Homocarnosine | 31.4 ± 8.12 (n=11) | 39.7 ± 7.55 (n=10) | |
| | Anserine | 866 ± 310 (n=11) | 1523 ± 350 (n=10) | |
| | Balenine | n.d. | 0.93 ± 0.43 (n=10) | |
| | N-acetylcarnosine | n.d. | 1.84 ± 0.25 (n=10) | |
| Vastus Lateralis | Carnosine | NA | NA | 2367 ± 1048 (n=8) |
| | Homocarnosine | | | 7.57 ± 2.09 (n=8) |
| | Anserine | | | 34.6 ± 14.6 (n=8) |
| | Balenine | | | 5.94 ± 3.57 (n=8) |
| | N-acetylcarnosine | | | 16.4 ± 9.91 (n=8) |
| Pectoralis Major | Carnosine | NA | NA | 1849 ± 1102 (n=10) |
| | Homocarnosine | | | 15.7 ± 10.3 (n=10) |
| | Anserine | | | 46.5 ± 27.9 (n=10) |
| | Balenine | | | 5.44 ± 4.70 (n=10) |
| | N-acetylcarnosine | | | 34.5 ± 58.8 (n=10) |
| Diaphragm | Carnosine | 9.18 ± 10.8 (n=4) | 56.2 ± 18.9 (n=10) | NA |
| | Homocarnosine | n.d. | n.d. | |
| | Anserine | 58.0 ± 25.0 (n=4) | 106 ± 21.5 (n=10) | |
| | Balenine | n.d. | n.d. | |
| | N-acetylcarnosine | n.d. | n.d. | |
| Heart [1] | Carnosine | 13.6 ± 6.39 (n=11) | 27.4 ± 6.54 (n=10) | 26.4 ± 20.9 (n=12) |
| | Homocarnosine | n.d. | n.d. | n.d. |
| | Anserine | 4.79 ± 2.23 (n=11) | 14.0 ± 7.83 (n=10) | 1.26 ± 1.55 (n=8/12) |
| | Balenine | n.d. | n.d. | n.d. |
| | N-acetylcarnosine | n.d. | n.d. | n.d. |
| **Central nervous system** | | **Mouse** | **Rat** | **Human** |
| Olfactory bulb | Carnosine | 1215 ± 448 (n=11) | 1207 ± 217 (n=10) | 79.3 ± 33.5 (n=3) |
| | Homocarnosine | 199 ± 44.7 (n=11) | 71.7 ± 12.4 (n=10) | 269 ± 54.1 (n=3) |
| | Anserine | 16.3 ± 3.63 (n=11) | 27.1 ± 17.0 (n=10) | 3.04 ± 2.64 (n=2/3) |
| | Balenine | n.d. | 2.11 ± 0.35 (n=10) | 1.24 ± 1.08 (n=2/3) |
| | N-acetylcarnosine | n.d. | 1.35 ± 0.11 (n=10) | 0.63 ± 0.55 (n=2/3) |
| Spinal cord (cervical) | Carnosine | 206 ± 12.0 (n=11) | 9.41 ± 4.62 (n=10) | 7.67 ± 4.91 (n=3) |
| | Homocarnosine | 777 ± 63.5 (n=11) | 122 ± 11.4 (n=10) | 491 ± 123 (n=3) |
| | Anserine | 14.0 ± 2.18 (n=11) | 7.61 ± 2.58 (n=10) | 2.83 ± 2.45 (n=2/3) |
| | Balenine | n.d. | 1.77 ± 0.68 (n=9/10) | n.d. |
| | N-acetylcarnosine | n.d. | n.d. | n.d. |
| Medulla oblongata | Carnosine | 129 ± 12.2 (n=5) | 11.0 ± 6.23 (n=7) | 3.40 ± 2.25 (n=3) |
| | Homocarnosine | 279 ± 24.9 (n=5) | 96.5 ± 11.8 (n=7) | 473 ± 131 (n=3) |
| | Anserine | 10.0 ± 1.29 (n=5) | 12.6 ± 14.5 (n=7) | 2.82 ± 2.44 (n=2/3) |
| | Balenine | n.d. | n.d. | n.d. |
| | N-acetylcarnosine | 1.37 ± 0.28 (n=5) | n.d. | n.d. |
| Frontal cortex [2] | Carnosine | 63.0 ± 18.9 (n=12) | 6.29 ± 2.60 (n=10) | 1.21 ± 0.79 (n=3) |
| | Homocarnosine | 197 ± 65.8 (n=12) | 70.1 ± 9.80 (n=10) | 312 ± 152 (n=3) |
| | Anserine | 5.30 ± 0.68 (n=12) | 4.58 ± 0.37 (n=10) | n.d. |
| | Balenine | n.d. | n.d. | n.d. |
| | N-acetylcarnosine | n.d. | n.d. | n.d. |
| Cerebellum | Carnosine | 86.5 ± 7.77 (n=11) | 4.64 ± 2.09 (n=10) | 0.66 ± 0.65 (n=2/3) |
| | Homocarnosine | 590 ± 83.5 (n=11) | 101 ± 16.4 (n=10) | 322 ± 114 (n=3) |
| | Anserine | 5.01 ± 0.76 (n=11) | 5.03 ± 0.79 (n=10) | 2.81 ± 2.44 (n=2/3) |
| | Balenine | n.d. | n.d. | n.d. |
| | N-acetylcarnosine | n.d. | n.d. | n.d. |
| Thalamus | Carnosine | 70.0 ± 13.3 (n=5) | 4.23 ± 1.33 (n=7) | 3.70 ± 3.27 (n=3) |
| | Homocarnosine | 169 ± 41.6 (n=5) | 76.4 ± 11.4 (n=7) | 577 ± 357 (n=3) |

| | | | | |
|---|---|---|---|---|
| | Anserine | 6.40 ± 0.55 (n=5) | 2.11 ± 0.25 (n=7) | 4.20 ± 0.12 (n=3) |
| | Balenine | n.d. | n.d. | n.d. |
| | N-acetylcarnosine | 1.22 ± 0.17 (n=5) | n.d. | n.d. |
| White matter [3] | Carnosine | 69.6 ± 12.2 (n=4) | 7.46 ± 1.41 (n=7) | 2.56 ± 1.77 (n=3) |
| | Homocarnosine | 155 ± 38.0 (n=4) | 100 ± 12.6 (n=7) | 219 ± 65.2 (n=3) |
| | Anserine | 6.71 ± 0.59 (n=4) | 2.18 ± 0.28 (n=7) | n.d. |
| | Balenine | n.d. | n.d. | n.d. |
| | N-acetylcarnosine | 1.01 ± 0.03 (n=4) | n.d. | n.d. |
| Hippocampus | Carnosine | 88.9 ± 65.0 (n=16) | 8.16 ± 4.61 (n=10) | NA |
| | Homocarnosine | 342 ± 156 (n=16) | 97.8 ± 21.1 (n=10) | |
| | Anserine | 17.2 ± 45.4 (n=16) | 4.61 ± 0.40 (n=10) | |
| | Balenine | n.d. | 1.62 ± 0.04 (n=10) | |
| | N-acetylcarnosine | n.d. | n.d. | |
| **Gastrointestinal system** | | **Mouse** | **Rat** | **Human** |
| Stomach wall | Carnosine | 0.99 ± 1.27 (n=3/4) | 2.69 ± 2.11 (n=10) | NA |
| | Homocarnosine | n.d. | n.d. | |
| | Anserine | n.d. | 1.86 ± 2.77 (n=8/10) | |
| | Balenine | n.d. | n.d. | |
| | N-acetylcarnosine | n.d. | n.d. | |
| Liver | Carnosine | 0.81 ± 0.49 (n=10/11) | 1.51 ± 0.47 (n=10) | 0.57 ± 0.54 (n=3/4) |
| | Homocarnosine | n.d. | n.d. | n.d. |
| | Anserine | n.d. | 1.84 ± 0.50 (n=10) | n.d. |
| | Balenine | n.d. | n.d. | n.d. |
| | N-acetylcarnosine | n.d. | n.d. | n.d. |
| Gallbladder | Carnosine | n.d. | NA | NA |
| | Homocarnosine | n.d. | | |
| | Anserine | 1.52 ± 0.70 (n=4) | | |
| | Balenine | n.d. | | |
| | N-acetylcarnosine | n.d. | | |
| Pancreas | Carnosine | 0.89 ± 0.38 (n=4) | 2.52 ± 1.53 (n=10) | NA |
| | Homocarnosine | n.d. | n.d. | |
| | Anserine | 0.76 ± 0.60 (n=3/4) | 1.93 ± 1.77 (n=8/10) | |
| | Balenine | n.d. | n.d. | |
| | N-acetylcarnosine | n.d. | n.d. | |
| Small intestine | Carnosine | 1.24 ± 0.99 (n=3/4) | 4.43 ± 4.85 (n=10) | NA |
| | Homocarnosine | n.d. | n.d. | |
| | Anserine | 1.87 ± 0.37 (n=4) | 5.49 ± 9.78 (n=7/10) | |
| | Balenine | n.d. | n.d. | |
| | N-acetylcarnosine | n.d. | n.d. | |
| Colon | Carnosine | 0.70 ± 0.55 (n=3/4) | 6.71 ± 5.77 (n=10) | NA |
| | Homocarnosine | n.d. | n.d. | |
| | Anserine | 1.41 ± 0.35 (n=4) | 6.01 ± 5.15 (n=9/10) | |
| | Balenine | n.d. | n.d. | |
| | N-acetylcarnosine | n.d. | n.d. | |
| **Adipose tissue** | | **Mouse** | **Rat** | **Human** |
| Adipose tissue (unspecified) | Carnosine | 7.45 ± 12.2 (n=10/11) | 4.61 ± 3.78 (n=10) | NA |
| | Homocarnosine | n.d. | n.d. | |
| | Anserine | 5.27 ± 5.97 (n=10/11) | 3.33 ± 2.12 (n=10) | |
| | Balenine | n.d. | n.d. | |
| | N-acetylcarnosine | n.d. | n.d. | |
| Adipose tissue (visceral) | Carnosine | NA | NA | 2.45 ± 1.80 (n=9/11) |
| | Homocarnosine | | | n.d. |
| | Anserine | | | n.d. |
| | Balenine | | | n.d. |
| | N-acetylcarnosine | | | n.d. |
| Adipose tissue (subcutaneous) | Carnosine | NA | NA | n.d. |
| | Homocarnosine | | | n.d. |
| | Anserine | | | n.d. |
| | Balenine | | | n.d. |
| | N-acetylcarnosine | | | n.d. |
| **Immune system** | | **Mouse** | **Rat** | **Human** |
| Thymus | Carnosine | 1.01 ± 0.27 (n=4) | n.d. | NA |

| | | | | |
|---|---|---|---|---|
| | Homocarnosine | n.d. | n.d. | |
| | Anserine | 0.95 ± 0.09 (n=4) | 1.23 ± 1.19 (n=6/10) | |
| | Balenine | n.d. | n.d. | |
| | N-acetylcarnosine | n.d. | n.d. | |
| Spleen | Carnosine | 2.45 ± 1.28 (n=11) | 3.52 ± 1.34 (n=10) | NA |
| | Homocarnosine | n.d. | n.d. | |
| | Anserine | 2.67 ± 1.82 (n=11) | 3.66 ± 1.72 (n=10) | |
| | Balenine | n.d. | n.d. | |
| | N-acetylcarnosine | n.d. | n.d. | |
| **Excretory system** | | **Mouse** | **Rat** | **Human** |
| Kidney | Carnosine | 16.3 ± 7.88 (n=11) | 2.35 ± 0.99 (n=10) | NA |
| | Homocarnosine | 0.63 ± 0.51 (n=7/11) | n.d. | |
| | Anserine | 7.38 ± 2.78 (n=11) | 2.63 ± 0.83 (n=10) | |
| | Balenine | n.d. | n.d. | |
| | N-acetylcarnosine | n.d. | 1.28 ± 0.14 (n=10) | |
| Kidney (medulla) | Carnosine | NA | NA | 1.62 ± 0.75 (n=5) |
| | Homocarnosine | | | 1.09 ± 1.00 (n=3/5) |
| | Anserine | | | n.d. |
| | Balenine | | | n.d. |
| | N-acetylcarnosine | | | n.d. |
| Kidney (cortex) | Carnosine | NA | NA | 2.31 ± 0.96 (n=5) |
| | Homocarnosine | | | n.d. |
| | Anserine | | | 1.99 ± 2.33 (n=4/5) |
| | Balenine | | | n.d. |
| | N-acetylcarnosine | | | n.d. |
| **Other** | | **Mouse** | **Rat** | **Human** |
| Eye | Carnosine | 6.40 ± 1.35 (n=4) | 17.0 ± 13.6 (n=10) | NA |
| | Homocarnosine | 1.49 ± 0.08 (n=4) | n.d. | |
| | Anserine | 5.25 ± 0.76 (n=4) | 5.58 ± 3.70 (n=10) | |
| | Balenine | n.d. | n.d. | |
| | N-acetylcarnosine | n.d. | n.d. | |
| Lung | Carnosine | 3.02 ± 1.84 (n=11) | 1.88 ± 1.12 (n=10) | 1.72 ± 1.14 (n=11/12) |
| | Homocarnosine | n.d. | n.d. | n.d. |
| | Anserine | 1.93 ± 1.47 (n=10/11) | 2.04 ± 1.96 (n=8/10) | n.d. |
| | Balenine | n.d. | n.d. | n.d. |
| | N-acetylcarnosine | n.d. | n.d. | n.d. |
| **Circulation & body fluids** | | **Mouse** | **Rat** | **Human** |
| Plasma | Carnosine | 1533 ± 503 (n=14) | 3944 ± 1262 (n=10) | 47.2 ± 97.7 (n=80/87) |
| | Homocarnosine | 114 ± 27.4 (n=14) | 120 ± 41 (n=10) | 66.4 ± 29.3 (n=87) |
| | Anserine | 1382 ± 720 (n=14) | 3351 ± 1437 (n=10) | 12.0 ± 17.9 (n=45/87) |
| | Balenine | n.d. | n.d. | 42.7 ± 101 (n=48/87) |
| | N-acetylcarnosine | 11.5 ± 7.2 (n=11/14) | 100 ± 25.1 (n=10) | 134.9 ± 109 (n=87) |
| Red blood cells | Carnosine | 25.8 ± 21.3 (n=8) | 43.4 ± 40.3 (n=10) | 99.4 ± 72.5 (n=7) |
| | Homocarnosine | n.d. | n.d. | 6.72 ± 6.33 (n=4/7) |
| | Anserine | 25.8 ± 26.2 (n=7/8) | 36.7 ± 26.6 (n=10) | 82.9 ± 62.6 (n=7) |
| | Balenine | n.d. | n.d. | 10.4 ± 7.00 (n=6/7) |
| | N-acetylcarnosine | 10.7 ± 0.39 (n=8) | 10.8 ± 0.19 (n=10) | 12.3 ± 1.32 (n=7) |
| Cerebrospinal fluid | Carnosine | NA | NA | 3.26 ± 2.89 (n=7/12) |
| | Homocarnosine | | | 2355 ± 1201 (n=12) |
| | Anserine | | | n.d. |
| | Balenine | | | n.d. |
| | N-acetylcarnosine | | | 17.9 ± 7.76 (n=12) |

Quantification of histidine-containing dipeptides (HCDs) using UHPLC-MS/MS. Concentrations are depicted in µmol/kg (tissue) or nM (fluids and red blood cells). Data are mean ± SD. NA, not applicable (not measured); n.d., not detectable (also if > 50% of samples had levels below the limit of detection of the respective HCD). The numbers between brackets indicate in how many samples the HCD was detected/measured. [1] Heart: unspecified region containing atrium and ventricle tissue (mouse, rat), right atrial appendage (human). [2] Frontal cortex: unspecified region (mouse, rat), superior frontal gyrus (human). [3] White matter: corpus callosum (mouse, rat), unspecified region (human).
